# Transition to self-compatibility associated with dominant *S*-allele in a diploid Siberian progenitor of allotetraploid *Arabidopsis kamchatica* revealed by *Arabidopsis lyrata* genomes

**DOI:** 10.1101/2022.06.24.497443

**Authors:** Uliana K. Kolesnikova, Alison Dawn Scott, Jozefien D. Van de Velde, Robin Burns, Nikita P. Tikhomirov, Ursula Pfordt, Andrew C. Clarke, Levi Yant, Alexey P. Seregin, Xavier Vekemans, Stefan Laurent, Polina Yu. Novikova

**Affiliations:** Department of Chromosome Biology, Max Planck Institute for Plant Breeding Research, Carl-von-Linne-Weg 10, Cologne, 50829, Germany; Department of Plant Sciences, Downing Street, University of Cambridge, Cambridge CB2 3EA, UK; Faculty of Biology, Lomonosov Moscow State University, Leninskie Gory Str. 1–12, Moscow, 119991, Russia; Papanin Institute for Biology of Inland Waters, Russian Academy of Sciences, Borok, Yaroslavl region, 152742, Russia; Future Food Beacon of Excellence and School of Biosciences, University of Nottingham, Sutton Bonington, LE12 5RD, UK; Future Food Beacon of Excellence and School of Life Sciences, University of Nottingham, Nottingham NG7 2RD, UK; Herbarium (MW), Faculty of Biology, M. V. Lomonosov Moscow State University, Moscow, 119991, Russia; Univ. Lille, CNRS, UMR 8198 – Evo-Eco-Paleo, F-59000 Lille, France; Department of Comparative Development and Genetics, Max Planck Institute for Plant Breeding Research, Carl-von-Linne-Weg 10, Cologne, 50829, Germany

## Abstract

A transition to selfing can be beneficial when mating partners are scarce, for example, due to ploidy changes or at species range edges. Here we explain how self-compatibility evolved in diploid Siberian *Arabidopsis lyrata,* and how it contributed to the establishment of allotetraploid *A. kamchatica*. First, we provide chromosome-level genome assemblies for two self-fertilizing diploid *A. lyrata* accessions, one from North America and one from Siberia, including a fully assembled S-locus for the latter. We then propose a sequence of events leading to the loss of self-incompatibility in Siberian *A. lyrata,* date this independent transition to ∼90 Kya, and infer evolutionary relationships between Siberian and North American *A. lyrata,* showing an independent transition to selfing in Siberia. Finally, we provide evidence that this selfing Siberian *A. lyrata* lineage contributed to the formation of the allotetraploid *A. kamchatica* and propose that the selfing of the latter is mediated by the loss-of-function mutation in a dominant *S*-allele inherited from *A. lyrata*.

## Introduction

Most angiosperms are hermaphroditic, with bisexual flowers producing both female and male gametes, and can thus potentially self-fertilize. Diverse self-recognition systems based on pollen–pistil interactions evolved repeatedly (Charlesworth et al. 2005; Zhao et al. 2022), preventing inbreeding, and subsequently, several independent transitions from outcrossing to self-pollination have occurred through degradation of these recognition systems (Shimizu and Tsuchimatsu 2015). A transition to selfing provides an immediate advantage in the face of low population density, often at the edges of the species distribution (Levin 2012). Pinpointing the genetic changes undermining self-rejection in nature not only improves our understanding of self-incompatibility mechanisms but also provides a more complete evolutionary history of the self-compatible species, providing essential context to understand their genome evolution (Guo et al. 2009; Slotte et al. 2013; Vekemans et al. 2014; Durvasula et al. 2017; Mable et al. 2017; Fulgione et al. 2018; Mattila et al. 2020).

In Brassicaceae, the sporophytic self-incompatibility (SI) system involves a self-pollen recognition mechanism determined by the *S*-locus, where two main genes are linked: the male *SCR* gene is expressed in tapetum cells of anthers, the protein is embedded into the pollen coat and serves as a ligand for the receptor kinase coded by the female *SRK* gene, which is expressed on the surface of the stigma (Stein et al. 1991; Schopfer et al. 1999; Takayama et al. 2000; Takayama et al. 2001; Takayama and Isogai 2005; Nasrallah 2019). A breakdown of SI and transition to self-compatibility occurs when recognition between *SCR* and *SRK* (or downstream signaling) leading to pollen rejection is impaired (Uyenoyama et al. 2001; Shimizu and Tsuchimatsu 2015; Mable et al. 2017). In outcrossing *Arabidopsis* species (e.g. *A. lyrata*, *A. halleri*, *A. arenosa*) more than 10 different S*-*haplotypes can segregate in a population (Castric and Vekemans 2004; Castric et al. 2008). This haplotypic diversity is essential for an SI system to function and has been maintained by frequency-dependent balancing selection for over 8 My (Mable et al. 2003; Castric and Vekemans 2004; Mable et al. 2004; Castric et al. 2008; Llaurens et al. 2008; Le Veve et al. 2022). A diploid outcrossing individual can possess two different *S*-alleles but often only one of them is expressed due to dominance, thus increasing the chances of reproduction (Hatakeyama et al. 2001; Kusaba et al. 2002; Prigoda et al. 2005; Okamoto et al. 2007), although co-dominance has also been reported (Prigoda et al. 2005; Llaurens et al. 2008). Expression of only one *S*-allele increases the chances for successful mating in heterozygous outcrossers, however, which of the *S*-alleles will be expressed can differ in pollen and stigma (Bateman 1954). Pollen-driven dominance is more thoroughly described and is conditioned by different trans-acting microRNA precursors and their targets on recessive *S*-alleles. MicroRNAs produced by dominant *S*-alleles silence the expression of the *SCR* gene on recessive *S*-allele through methylation of a 5’ promoter of *SCR* (Kusaba et al. 2002; Shiba et al. 2006); (Tarutani et al. 2010; Durand et al. 2014; Fujii and Takayama 2018). As dominance is uncoupled from self-recognition in this system, a dominant loss-of-function mutation is possible and would yield a self-compatible phenotype in a heterozygous individual. The ancestral state in the genus *Arabidopsis* is outcrossing due to self-incompatibility. However, self-compatible species have evolved multiple times: in the model species *A. thaliana*, and allotetraploids *A. suecica* and *A. kamchatica*. One of the early challenges for a new polyploid is the scarcity of compatible karyotypes for mating, and competition with established nearby diploids (Levin 1975). Selfing alleviates such challenges. In *A. suecica,* the transition to self-compatibility was likely immediate following the cross between an *A. thaliana* with a non-functional dominant S-haplotype (Tsuchimatsu et al. 2010) and an outcrossing *A. arenosa* (Novikova et al. 2017). However, the origin of self-compatibility in *A. kamchatica* is less clear, as the species originated from multiple crosses between *A. lyrata* and *A. halleri* in East Asia (Shimizu et al. 2005; Shimizu-Inatsugi et al. 2009; Tsuchimatsu et al. 2012; Paape et al. 2018). While *A. halleri* is an obligate outcrosser, *A. lyrata* is predominantly self-incompatible with described self-compatible populations restricted to the Great Lakes region of North America (Mable et al. 2005; Foxe et al. 2010; Willi and Määttänen 2010; Griffin and Willi 2014) from which subarctic and arctic selfing *A. arenicola* in Canada and Greenland may have originated. (Willi et al. 2022). A selfing individual of *A. lyrata* collected in Yakutia has been reported as genetically closest to the *A. lyrata* subgenome of *A. kamchatica* (Shimizu-Inatsugi et al. 2009; Paape et al. 2018), but the evolutionary history of this selfing lineage and *S*-locus genotype have not been described.

Here we ask (1) how and when self-compatibility evolved and spread in Siberian *A. lyrata*. (2) Is it plausible that *A. lyrata* was already self-compatible when it contributed to allopolyploid *A. kamchatica*? (3) Could a loss of self-incompatibility in only one of the diploid ancestors (*A. lyrata)* be sufficient to render *A. kamchatica* self-compatible? Broad sampling combining live and herbarium collections allowed us to describe the selfing lineage of *A. lyrata* in Siberia ranging between Lake Taymyr and Chukotka, across north-central and eastern Russia. We first present chromosome-level assemblies of a Siberian selfing *A. lyrata* and the reference North American selfing accession (Hu et al. 2011), characterize the genomic and structural differences between them and describe the *S*-locus structure and the likely mechanism of the failure of self-incompatibility in the Siberian selfing populations. Using demographic modeling we date the transition to selfing in Siberian *A. lyrata* and suggest that it happened prior or concurrent with the formation of allopolyploid *A. kamchatica*. We confirm that the Siberian selfing *A. lyrata* was likely one of the progenitors of the allotetraploid *A. kamchatica* using overall genetic relatedness assessment and the phylogeny of the *SRK* gene in the *S*-locus. Together, our results suggest that one of the allopolyploid *A. kamchatica* origins and its transition to selfing was facilitated by the loss-of-function in the dominant *S*-allele inherited from Siberian *A. lyrata*.

## Results

### Genome assembly of the selfing Siberian NT1 accession

We grew seeds of *A. lyrata* collected from three populations in Yakutia (Supplementary Table 1, (Supplementary Fig. 1A) in the greenhouse (see Methods) and noticed that plants from NT1 population formed long fruits (Supplementary Fig. 1B), suggesting self-compatibility. We observed that flowers of the selfing NT1 accession appeared to be smaller compared to flowers of outcrossing plants and another selfing accession MN47 from North America (Supplementary Fig 1C-E). We confirmed that self-pollen successfully germinated in a selfed NT1 accession and made pollen tubes, whereas self-pollination of an outcrossing plant from the NT8 population did not result in pollen tube growth (Supplementary Figure 2).

We extracted HMW DNA from NT1 leaf tissue and obtained 1,100,878 high-fidelity (HiFi) PacBio reads with N50 read length of 14,161bp (total length of raw read sequences is ∼15,9Gbp). We assembled those reads using the Hifiasm (Cheng et al. 2021) into 1,070 contigs with N50 of 5.508 MB. We scaffolded these contigs further along the MN47 *A. lyrata* assembly (Hu et al. 2011) with RagTag (Alonge et al. 2019) reaching chromosome-level with a scaffold N50 of 24.641Mb. We then assessed the completeness of the NT1 *A. lyrata* genome assembly using BUSCO and found 4,463 complete and single-copy (97.1%), 88 complete and duplicated (1.9%), 7 fragmented (0.2%), and 38 missing genes (0.8%) from the Brassicales_odb10 set. Repeated sequences composed about 49.9% of the assembly. We annotated 28,596 genes by transferring gene annotation from the reference *A. lyrata* genome (Rawat et al. 2015) using Liftoff (Shumate and Salzberg 2020).

Various papers (Long et al. 2013; Slotte et al. 2013; Henry et al. 2014; Burns et al. 2021; Dukić and Bomblies 2022) have reported potential artifacts in the reference *A. lyrata* MN47 (version 1 or v1) genome assembly (Hu et al. 2011). Our comparison of the Siberian NT1 with the MN47 v1 *A. lyrata* reference genome indicated multiple structural variants in the same genomic regions as those between the genomes of MN47 v1 and the *A. arenosa* subgenome of *A. suecica* (Fig.1A), MN47 v1 and *Capsella rubella,* and MN47 v1 and a diploid *A. arenosa* (Supplementary Table 2) (Long et al. 2013; Slotte et al. 2013; Burns et al. 2021; Dukić and Bomblies 2022). We confirmed the existence of such artifacts and corrected them through long-read DNA sequencing (Supplementary Table 2). Specifically, we obtained 868,563 HiFi reads of the MN47 accession with N50 length of 20,206 bp (total length of raw read sequences is ∼17,6Gbp; ∼80x coverage). In total we assembled ∼244Mb in 820 contigs with an N50 of 23.506Mb, indicating that full chromosome arms of MN47 were assembled as single contigs. Contigs were scaffolded into eight chromosomes using the genomes of MN47 v1 and NT1 as guides. The scaffolded contigs amount to ∼209Mb. Completeness of the new MN47 v2 *A. lyrata* genome assembly by BUSCO was 4,544 complete and single-copy (97.1%), 83 complete and duplicated (1.8%), 8 fragmented (0.2%), and 44 missing genes (0.9%) from the Brassicales_odb10 set. The placement and orientation of contigs in the scaffolds were corrected using previously published Hi-C data (Zhu et al. 2017) and by manual examination of the long reads (see *Methods*, Supplementary Figs. 3-7).

**Figure 1.**
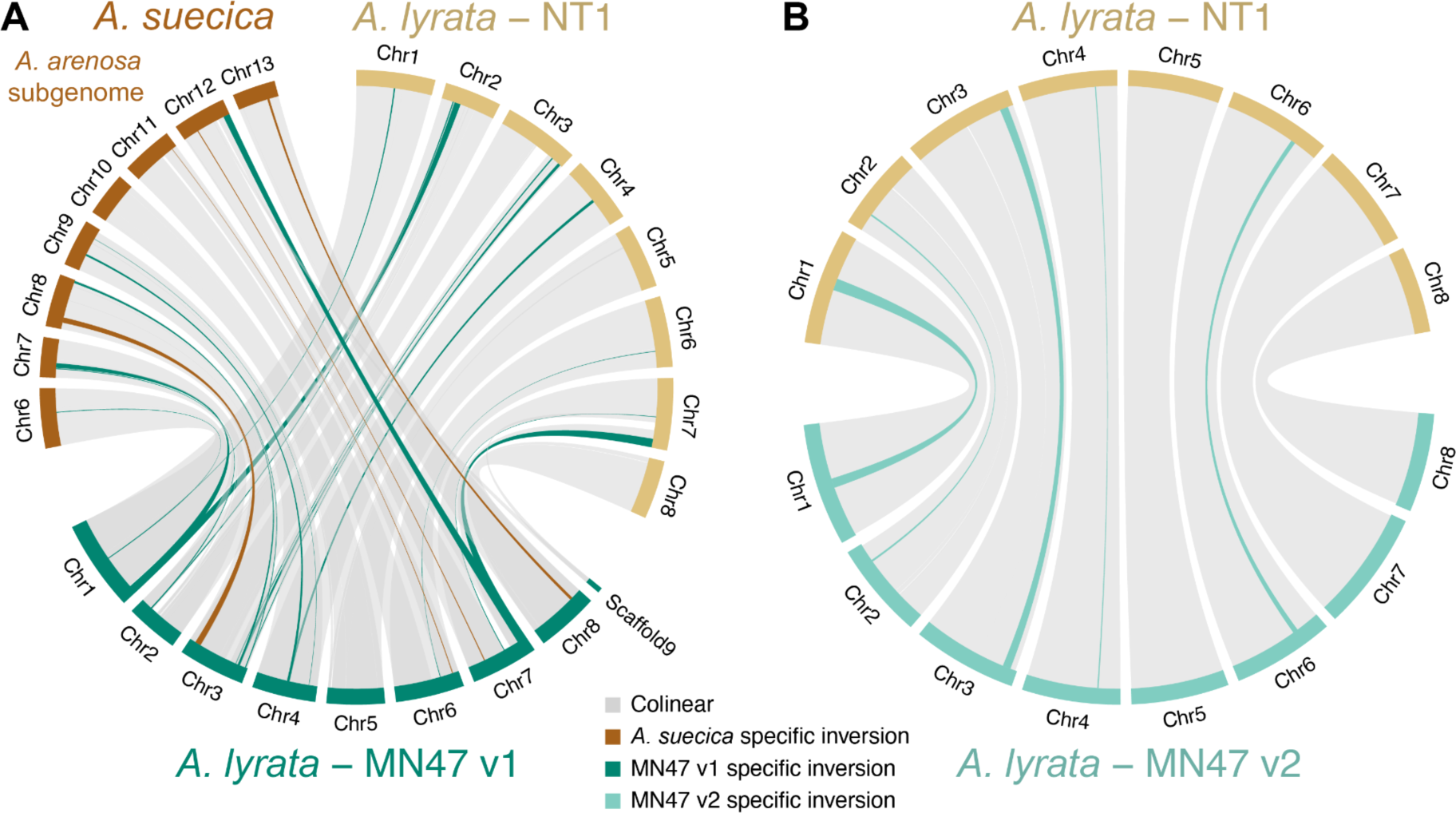
Segregating large structural variants in *A. lyrata*. (A) Eleven large inversions (dark green) between North American MN47 (dark green) and Siberian NT1 (dark yellow) *A. lyrata* are also observed between MN47 and the *A. arenosa* subgenome of *A. suecica* (brown), but are not observed in the comparison between NT1 and *A. suecica* (Supplementary Table 2), which suggests these inversions are likely artifacts in the original MN47 assembly. (B) Segregating inversions in *A. lyrata* observed following re-assembly by long reads and manual curation using Hi-C data of the North American MN47 genome (turquoise) and its alignment to the Siberian NT1 (dark yellow) *A. lyrata* genome. Five inversions that are unique to MN47 (the longest being ∼2.4 Mb in size) are highlighted in turquoise.

Our re-assembled long read-based MN47 v2 genome confirmed the existence of the expected false structural variants in the MN47 v1 genome (Figure 1, Supplementary Figures 3-7) which we were able to fix in the v2. The comparison of the MN47 v2 and NT1 genomes revealed several large inversions segregating in *A. lyrata*, the largest of which (∼2.4 Mb) is on chromosome 1. All the identified inversions between the genomes comparisons are listed in Supplementary Table 2. Inversions between *A. lyrata* MN47 and NT1 accessions are listed in Supplementary Table 1. In each genome comparison, we identified multiple alleles of structural variants at the end of chromosome 3. This may be explained by the fact that one of the nucleolar organizer regions (NORs) of *A. lyrata* is located at the end of chromosome 3 (Lysak et al. 2006). We confirmed that chromosome 3 contains a partially assembled NOR using BLAST. Overall, we have assembled high-quality chromosome-level genomes for two *A. lyrata* accessions and through pairwise genome alignment we identified several inversions up to 2.4 Mb long segregating in the species.

### Breakdown of the SI system in Siberian *A. lyrata* NT1

Both genes flanking the *S*-locus (*U-box* and *ARK3*) were assembled in a single contig in the HiFi assembly before any scaffolding, indicating the entire ∼44.5kb *S*-locus of the NT1 accession was fully assembled. We further confirmed the completeness of the *S*-locus by mapping PacBio reads back to the assembly, and found even coverage spanning the *S*-locus with no gaps (Supplementary Fig. 8). BLAST analysis of *SRK* and *SCR* sequences from the known S-haplotypes (Supplementary Data 1 and 2) (N. A. Boggs et al. 2009; Tsuchimatsu et al. 2010; Guo et al. 2011; Goubet et al. 2012; Tsuchimatsu et al. 2012) revealed no hits for *SRK*, and one hit for *SCR* from the *A. halleri* S12 haplogroup (Figure 2b). Due to long-term frequency-dependent balancing selection on the *S*-locus in Brassicaceae, relatedness among S-haplotypes is not consistent with species relatedness, such that the closest sequences to *A. halleri* S12 (AhS12) are not other *A. halleri* S-haplotypes but rather specific S-haplotypes from *A. lyrata* S42 (AlS42) and *A. kamchatica* D (Ak-D) (Wright 1939; Vekemans and Slatkin 1994; Mable et al. 2003; Castric and Vekemans 2004; Kamau and Charlesworth 2005; Castric et al. 2008; Llaurens et al. 2008; Tsuchimatsu et al. 2012; Roux et al. 2013). We estimated a phylogeny of the known SCR protein sequences (Guo et al. 2011; Goubet et al. 2012) and the manually annotated NT1 *A. lyrata SCR* sequence from the BLAST results (Figure 2a). As expected, the SCR phylogeny has a different topology than the species phylogeny, as S-haplotypes are trans-specifically shared across *Arabidopsis*. The SCR phylogeny confirms that the closest haplotype to the NT1 *A. lyrata S*-locus is the S12 haplotype from *A. halleri* (AhS12).

**Figure 2.**
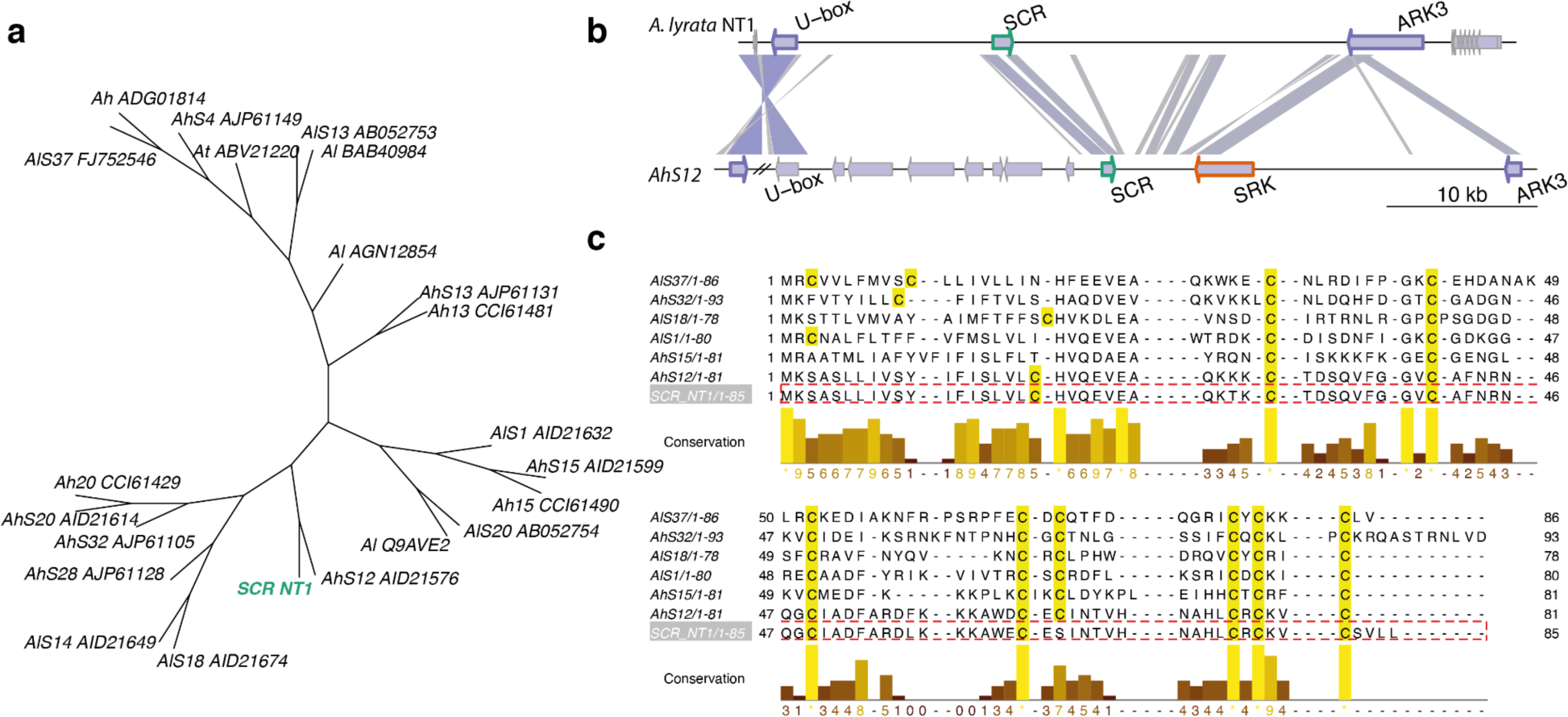
*S*-locus structure of the Siberian NT1 selfing *A. lyrata* population. (A) Phylogenetic tree of SCR proteins reveals clustering of NT1 SCR (green) and AhS12. (B) Comparison of the *S*-locus region of the *A. lyrata* NT1 genome assembly with the *A. halleri* S12 haplotype (Durand et al. 2014). Links between *S*-loci are colored according to the BLAST scores from highest (blue) to lowest (gray). *SCR*, *SRK* and flanking *U-box* and *ARK3* genes have green, orange and purple borders, respectively. *SRK* gene appears to be completely absent from the *S*-locus of the NT1 *A. lyrata* selfing accession. The only BLAST hit to *SRK* is a spurious hit to *ARK3* as they both encode receptor-like serine/threonine kinases. (C) Protein sequence alignment of *S*-locus *SCR* genes from *A. halleri* and *A. lyrata*, including NT1. One of the eight conserved cysteines important for structural integrity has been lost from the NT1 SCR protein.

We compared the structures of the AhS12 and NT1 *S*-loci (Figure 2b) and confirmed the absence of *SRK*, (i.e. the female component of the self-incompatibility system) which is sufficient to explain the selfing nature of the NT1 accession. We also mapped short reads from NT1 to the NT1 genome assembly plus the intact AhS12 sequence from *A. halleri* containing *SRK*, and found no reads mapped to *SRK* (Supplementary Fig. 9C). This provides additional confirmation of a complete loss of *SRK* from the NT1 *S*-locus. Analyzing the SCR protein sequences more closely, we also observed a loss of one of the eight conserved cysteines in the NT1 SCR sequence, which are important in protein-folding and the recognition of the SCR ligand by the SRK receptor (Kusaba et al. 2001; Mishima et al. 2003; Tsuchimatsu et al. 2010) (Supplementary Fig. 10A). This suggests the SCR protein is non-functional in the NT1 *A. lyrata* accession. We tested for expression of the *SCR* gene in the flowers of NT1 using RNAseq and did not detect any transcript of the AhS12 *SCR* (Supplementary Fig. 9A,B), though this may be due to the timing of floral development as expression of *SCR* is transient (Burghgraeve et al. 2020). Sequence comparison of the *SCR* region between AhS12 and NT1 showed high similarity in the promoter region (Supplementary Fig. 9D) indicating structural re-arrangements did not cause loss of expression - but nucleotide substitutions at critical sites cannot be excluded. To verify whether SCR is indeed non-functional and/or not expressed in NT1, we performed controlled crosses, fertilizing an outcrossing *A. lyrata* accession (NT8.4-24, which has a functional AhS12 haplogroup) with NT1 pollen, resulting in successful pollen tube growth (Supplementary Figure 11). This outcome is possible if (1) the SCR protein from the NT1 accession could not be recognized by SRK receptors from the same AhS12 haplogroup or (2) the *SCR* gene was not expressed at all. Both of these scenarios lead to the conclusion that the SCR gene is non-functional in the NT1 selfing Siberian *A. lyrata* accession. There is, however unlikely, an additional possibility (3): a self-compatible reaction could be possible with a functional SCR in NT1 if the *SRK* gene from haplogroup AhS12 was not expressed in the outcrossing maternal plant (NT8.4-24). We describe scenario (3) as improbable because outcrossing maternal plant NT8.4-24 is heterozygous at the S-locus, possessing two *S*-alleles: AhS12 and AlS25. The latter is known to be either co-dominant or recessive to AhS12 as it belongs to a lower dominance class (Llaurens et al. 2008; Durand et al. 2014), therefore AhSRK12 is most likely expressed in NT8.4-24.

According to the classification of S-haplotypes, AhS12 belongs to dominance class IV (the most dominant class) and it is documented that it has an sRNA precursor which can silence the expression of *SCR* genes from S-haplotypes belonging to class I, II and III (Durand et al. 2014; Burghgraeve et al. 2020). Indeed, by BLAST analysis, we identified an sRNA precursor sequence in the NT1 *S*-locus assembly similar to the mirS3 precursor of *A. halleri* S12 haplotype (Durand et al. 2014), suggesting a conserved dominance mechanism of *A. lyrata* S12.

### Population-level re-sequencing confirms that selfing Siberian *A. lyrata* contributed to *A. kamchatica* origin

#### Sampling

We sequenced an additional nine *A. lyrata* accessions collected during the same expedition (Supplementary Table 1), 10 herbarium samples of *A. lyrata* from Taymyr, Yakutia, Kamchatka and Chukotka dating from 1958–2014, and 19 herbarium samples of *A. kamchatica* using the same Illumina NovaSeq platform (150bpPE) (see Methods and Supplementary Table 1). The herbarium samples were obtained from the Moscow University Herbarium (Seregin 2023). Our dataset also included previously published whole genome resequencing data from the diploid *A. lyrata* collected in the same region, allotetraploid *A. kamchatica* samples (Shimizu-Inatsugi et al. 2009; Novikova et al. 2016; Paape et al. 2018) (Supplementary Table 1, Figure 3A) and European *A. lyrata* samples (Takou et al. 2021) as outgroups (Supplementary Table 1).

**Figure 3.**
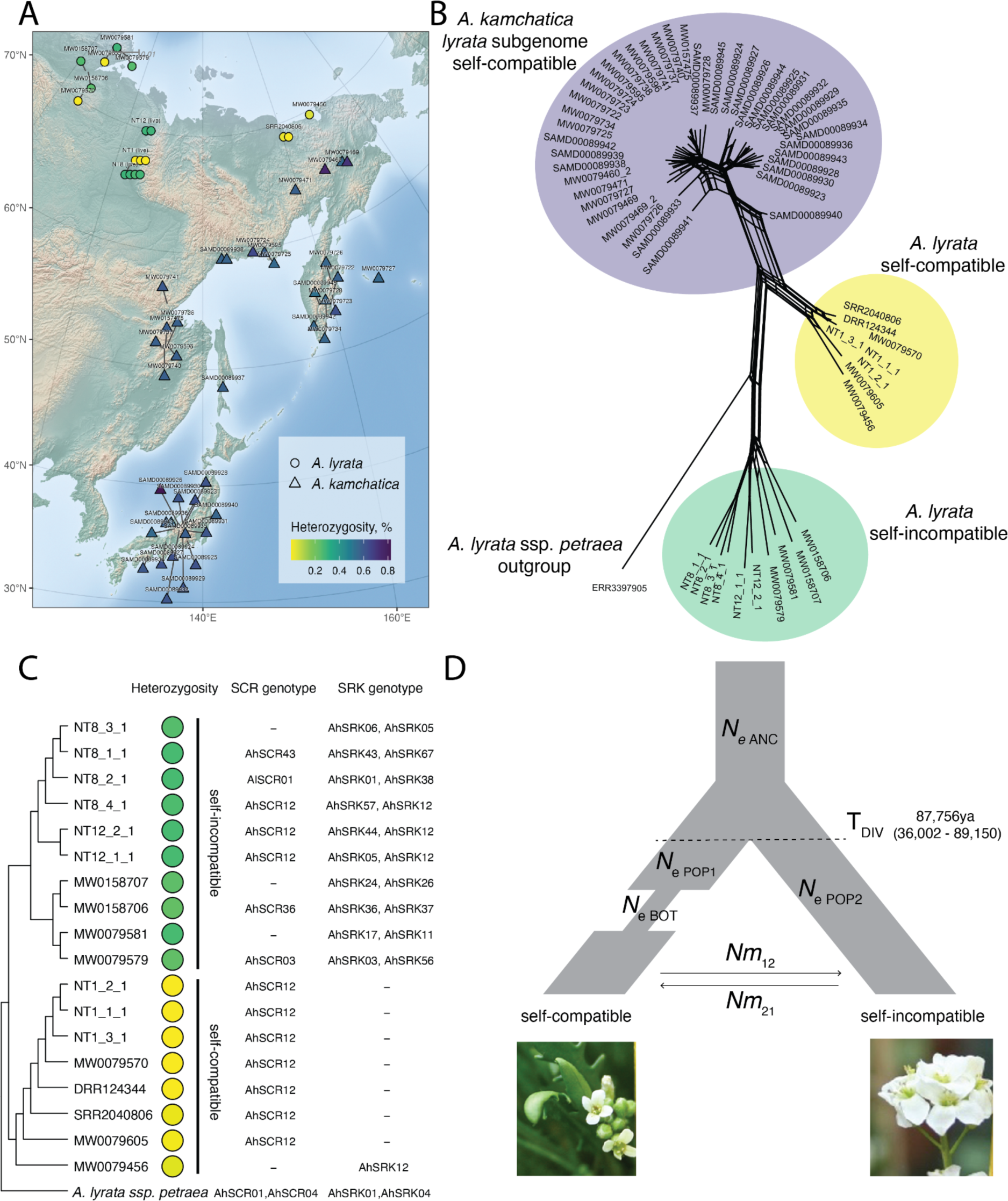
(A) Map of short-read sequenced Siberian *A. lyrata* (circles) and *A. kamchatica* (triangles). Live *A. lyrata* accessions names start with NT, herbarium sample names start with MW, a previously published sample of *A. lyrata* has been assigned the SRR and DRR prefix, and *A. kamchatica* samples start with SAMD. Colors indicate heterozygosity per sample, calculated by the percent of heterozygous sites. (B) Network depiction of Nei’s D between individuals shows that selfing *A. lyrata* is genetically closer to *A. kamchatica* than the outcrossing populations. While the network is drawn as unrooted, an outgroup accession provides context for interpretation. Individual genetic distances are also shown as heatmap in Supplementary Figure 14. (C) Neighbor-joining tree of Siberian *A. lyrata* accessions with heterozygosity and genotyped *SCR* snd *SRK* alleles. (D) Best-fit demographic model of divergence, a bottleneck in selfers, and asymmetric migration between selfing and outcrossing lineages, with parameter estimate for divergence time.. T_DIV_ = time of divergence between selfing and outcrossing lineage (origin of selfing), Ne_ANC_= effective population size of ancestor lineage, Ne_POP1_ = effective population size of selfing lineage, Ne_POP2_ = effective population size of outcrossing lineage, T_BOT_ = time of bottleneck in selfing lineage, Nm_12_ and Nm_21_ = the number of migrants between selfing and outcrossing lineages. Further values are reported in Table 1. Point estimates and confidence intervals are reported in Table 1; point estimates for all tested models are reported in Supplementary Table 4.

**Table 1.** Point estimates and 95% confidence intervals for the best-fit demographic model. Parameter estimates are reported in number of individuals and years.

|  | Parameter estimates |  |  |  |  |  |  |
| --- | --- | --- | --- | --- | --- | --- | --- |
|  | **N<sub>e</sub> POP1** | **N<sub>e</sub> POP2** | **N<sub>e</sub> ANC** | **T<sub>Div</sub>** | **N<sub>m</sub>12** | **N<sub>m</sub>21** | **N<sub>e</sub> BOT** |
| best point estimate | 526,490 | 33,400 | 129,243 | 87,756 | 2.69E-01 | 3.79E+00 | 14 |
| CI 2.5% | 5,602 | 20,224 | 1,418 | 36,002 | 7.61E-05 | 8.35E-06 | 5 |
| CI 97.5% | 552,758 | 56,554 | 275,637 | 89,150 | 1.37E+00 | 1.11E+01 | 48 |

#### Defining selfing *A. lyrata* by heterozygosity

To determine whether multiple selfing populations might exist in the examined geographic region, we first calculated the percent of heterozygous sites for each individual (Supplementary Table 1, Figure 3A) mapped to NT1 reference. Two modes on the heterozygosity levels were apparent in our *A. lyrata* dataset (Supplementary Fig. 12), which we assign as selfing (0.012% on average with 0.046% maximum value in Supplementary Table 1, indicated with yellow markers on Figure 3) and outcrossing (0.27% on average with 0.288% maximum value in Supplementary Table 1 within *A. lyrata* samples, indicated by green markers on Figure 3). This heterozygosity-based assignment is supported by our observations of individuals growing in the greenhouse: NT1 populations produced seeds without crosses, whereas NT8 and NT12 populations did not. Allotetraploid *A. kamchatica* co-occuring in the same geographical region is also self-compatible. To ensure that none of our *A. lyrata* samples were misclassified, we first mapped allotetraploid *A. kamchatica* samples in the same way to the NT1 *A. lyrata* reference without separating subgenomes. The majority of the SNPs in *A. kamchatica* represent divergent sites between the two subgenomes, which explains its high heterozygosity levels, clearly distinct from selfing *A. lyrata* samples (Supplementary Fig. 12 and Figure 3).

#### Genotyping S-alleles in outcrossers

We genotyped *S*-alleles of all the short-read sequenced accessions in our dataset by using a genotyping pipeline for *de novo* discovery of divergent alleles (Genete et al. 2020) with both *SCR* and *SRK* sequences as the reference allele databases (Schierup et al. 2001; Mable et al. 2003; Bechsgaard et al. 2004; Castric and Vekemans 2007; Castric et al. 2008; Nathan A. Boggs et al. 2009; Guo et al. 2009; Castric et al. 2010; Guo et al. 2011; Goubet et al. 2012; Dwyer et al. 2013; Durand et al. 2014; Mable et al. 2017; Tsuchimatsu et al. 2017; Mable et al. 2018; Takou et al. 2020; Kodera et al. 2021) (Supplementary Data 1 and 2, Supplementary Table 1). For each outcrossing individual we find two different *SRK* alleles and at most one *SCR* allele (Fig. 3C). Identifying *SCR* alleles is more difficult than *SRK*, likely due to an incomplete *SCR* database rather than these genes being absent in outcrossing individuals.

#### Selfing Siberian *A. lyrata* is fixed for AhS12

All the self-compatible (low-heterozygosity) *A. lyrata* samples shared the same *S*-haplogroup – AhS12 (Fig. 3C): either by *SCR* or *SRK* genotype. Most of the self-compatible accessions, with exception of MW0079456, did not have *SRK* genotype. As the *SRK* database is robust and we confirmed the absence of *SRK* from the full-length NT1 assembly, the lack of *SRK* genotypes in the self-compatible Siberian *A. lyrata* accessions is likely due to gene loss.

#### *S*-allele of the self-compatible Siberian *A. lyrata* matches the most common *A. lyrata*-inherited *S*-allele in *A. kamchatica*

In the pool of *A. kamchatica* samples we identified five *SRK* alleles (Supplementary Table 1) using the same genotyping pipeline (Genete et al. 2020), consistent with Tsuchimatsu et al. 2012, where AhS12 (AkS-D) and AhS02 (AkS-E) are shown to be *A. lyrata*-inherited, while AhS26 (AkS-A), AhS47 (AkS-B) and AhS1 (AkS-C) are *A. halleri*-inherited (Tsuchimatsu et al. 2012). *A. lyrata*-inherited AhS12 (AkS-D) is the most common *SRK* allele (43.67%) on the *A. lyrata* subgenome of *A. kamchatica* and matches the *S*-haplotype of the Siberian self-compatible *A. lyrata* lineage. The full *SRK* gene sequence from Siberian self-compatible *A. lyrata* accession MW0079456 forms a monophyletic group with *A. kamchatica SRK* sequences from the S12-haplogroup while outcrossing *A. lyrata SRK* sequences from the S12-haplogroup are in a distinct clade separate from *A. kamchatica* (Supplementary Fig. 13A, Supplementary Data 3).

#### *S*elf-compatible Siberian *A. lyrata* lineage is genetically closest to *A. kamchatica*

To estimate the relatedness between *A. lyrata* and *A. kamchatica* we analyzed only the *A. lyrata* subgenome of *A. kamchatica*. We split the subgenomes of *A. kamchatica* by mapping accessions simultaneously to NT1 *A. lyrata* and *A. halleri ssp. gemmifera* (Briskine et al. 2016) reference genomes, and used only the *A. lyrata* portion for further analysis. In addition to the SRK phylogeny, network analysis based on genetic distance (Nei’s D) (Nei 1972) between individuals for 4,141 biallelic SNPs at four-fold degenerate sites suggests an overall closer genome relatedness between Siberian selfing *A. lyrata* and *A. lyrata* subgenome of *A. kamchatica,* compared to Siberian outcrossing *A. lyrata* and the *A. lyrata* subgenome of *A. kamchatica* (Figure 3B). These individual pairwise genetic distances are further represented as a heatmap (Supplementary Figure 14). The same relationships are also shown in the maximum likelihood phylogeny based on the same SNP data, where selfing Siberian *A. lyrata* populations form a clade with *A. lyrata* subgenome of *A. kamchatica* (Supplementary Fig. 13B), while outcrossing Siberian *A. lyrata* is more distantly related. This is consistent with the previously published results showing that a selfing *A. lyrata* accession from Siberia (lyrpet4 -DRR124344) is genetically closest to *A. kamchatica* (Shimizu-Inatsugi et al. 2009; Paape et al. 2018).

#### Demographic modeling suggests that self-compatible Siberian *A. lyrata* lineage originated around 90kya

The observation that all the Siberian selfing *A. lyrata* accessions share the same *S-*haplotype suggests they may have originated from a single breakdown of self-incompatibility. The calculated total nucleotide diversity in 10kb windows for selfing *A. lyrata* has a mean value of 0.11% (95% CI [0.105 – 0.118]), which is about 7.5 times lower compared to 0.84% (95% CI [0.818 – 0.87]) in the outcrossing Siberian *A. lyrata* population. Though the selfing lineage in Siberia likely originated from a single founder, the joint allele frequency spectrum between selfing and outcrossing Siberian *A. lyrata* shows a considerable amount of shared polymorphism (genome-wide non-genic and excluding pericentromeric and centromeric regions - 54,772 SNPs shared, vs 128,393 SNPs private to the selfer lineage; Supplementary Figure 14). This may be because the founder was a heterozygous outcrosser and a certain amount of gene flow does occur between lineages, as self-compatibility does not prevent plants from mating with outcrossers.

To further investigate the relationships between selfing and outcrossing populations and to date the self-incompatibility breakdown, we implemented a series of demographic models in fastsimcoal26 (Excoffier et al. 2013). The best-fit model is shown (Fig. 3D), which includes divergence between selfers and outcrossers with a subsequent bottleneck in the selfing lineage, with asymmetric introgression between populations. The estimate of divergence time (TDIV) in this model is ∼90 ka (87,756), though we suggest caution when interpreting such estimates. All tested models can be viewed in Supplementary Figure 15, with corresponding parameters in Supplementary Table 4 and input files on GitHub (https://github.com/novikovalab/selfing_Alyrata).

#### *S*-allele dominance is retained in the self-compatible Siberian *A. lyrata* lineage

Above we show that the self-compatible Siberian *A. lyrata* lineage is fixed for AhS12, which belongs to a dominant class of *S*-alleles in *Arabidopsis* (Durand et al. 2014). To test whether dominance is retained in *A. lyrata* NT1 despite the loss of the self-recognition function in the AhS12 *S*-allele, we conducted two crosses with self-incompatible *A. lyrata* plants (TE10.3-2 and TE11) as maternal plants and NT1 as pollen donor. The resulting F1 plants had different combinations of *S*-alleles and were self-compatible in two combinations, where the maternally inherited *S*-allele from a self-incompatible plant was from a lower dominance class than AhS12 (Table 2, Figure 4A, B and Supplementary Fig. 16). F1 plants with AhS1/AhS12 and AhS63/AhS12 combinations of *S*-alleles are self-compatible, while F1 plants with AhSRK54/AhS12 combination are self-incompatible. AhS1 is recessive to AhS12 in *A. halleri* (Durand et al. 2014), and AhS63 belongs to class III of dominance (corresponding to AlS41 in Mable et al. 2018) which is expected to be recessive to the class IV AhS12 allele in *A. lyrata* (Prigoda et al. 2015).

**Figure 4.**
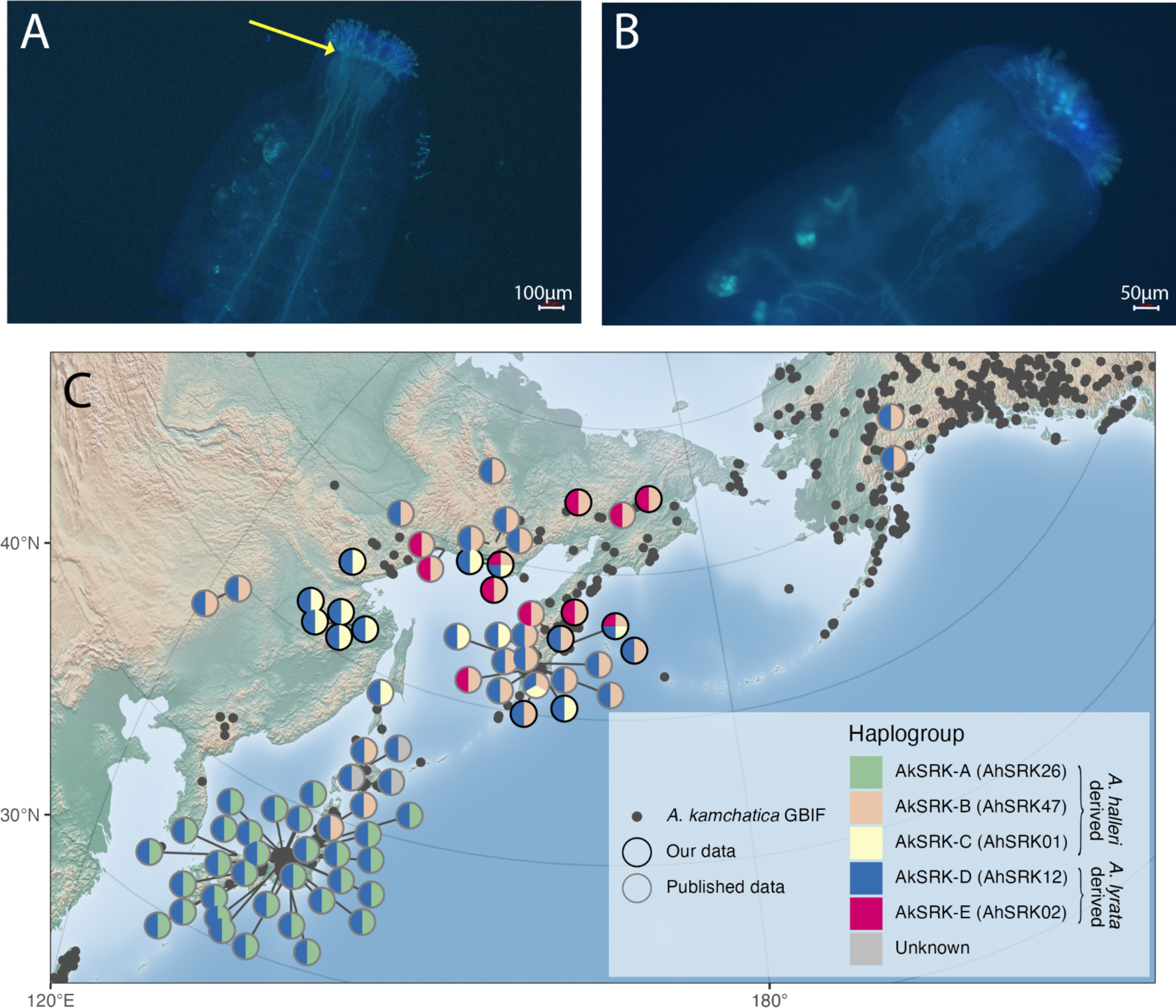
(A) Self-pollinated F1 progeny (F1.1-1) resulting from a cross between a self-incompatible (shown in B) **♀** TE10.3-2 *A. lyrata* accession and ♂ NT1 self-compatible *A. lyrata* accession shows pollen tube growth (yellow arrow) and dominance of self-compatibility in the F1 generation. (B) Self-pollinated self-incompatible *A. lyrata* accession TE10.3-2 (used as the maternal plant in A) shows no pollen tube growth, demonstrating its self-incompatibility. (C) The geographical distribution of *A. kamchatica* S-haplotypes shows a strong population structure across the species range. Circles are individual accessions, with S-haplogroups indicated by colors of pie slices. *A. halleri* orthologous S-haplogroups are mentioned in the parenthesis next to the *A. kamchatica* S-haplogroups (AkS-A-E). Circle outline indicates either previously published data (grey) or newly reported accessions (black). *A. kamchatica* occurrences from GBIF are indicated by transparent grey dots.

**Table 2.** The genotypes and phenotypes of outcrossing mother plants (TE10.3-2 and TE11.1-2) and F1 progeny from their pollination by NT1 self-compatible *A. lyrata* accession with AhS12 S-allele (*SCR* present, *SRK* lost -Figure 2B). Some of the *SCR* genotypes have missing data “-” due to the incomplete *SCR* database. Note that in the F1s AhSRK12 is missing due to gene loss in NT1 (Figure 2).

| Accession | SRK genotype | SCR genotype | Mating type |
| --- | --- | --- | --- |
| TE10.3-2 | AhSRK01, AhSRK63 | AhSCR01, - | SI |
| TE11.1-2 | AhSRK01, AhSRK54 | AhSCR01, - | SI |
| NT1 | - | AhSCR12 | SC |
| TE10.3-2♀ x NT1♂ F1.1-1 | AhSRK01 | AhSCR01, AhSCR12 | SC |
| TE10.3-2♀ x NT1♂ F1.1-2 | AhSRK63 | AhSCR12, - | SC |
| TE11.1-2♀ x NT1♂ F1.2-1 | AhSRK54 | AhSCR12, - | SI |
| TE11.1-2♀ x NT1♂ F1.2-2 | AhSRK54 | AhSCR12, - | SI |

#### Ancestral dominant *S*-allele AhS12 with lost self-recognition function could promote *A. kamchatica* establishment

Multiple crosses between different *A. lyrata* and *A. halleri* have contributed to allopolyploid *A. kamchatica* (Shimizu et al. 2005; Shimizu-Inatsugi et al. 2009; Tsuchimatsu et al. 2012; Paape et al. 2018). This is also apparent in the strong population structure of the *S*-allele combinations inherited from different parental lineages (Figure 4C). The most common *S*-allele in *A. kamchatica* on the *A. lyrata* subgenome is AhS12 (AkS-D), which is also fixed in the self-compatible Siberian *A. lyrata* lineage.

Moreover, F1 crosses (Figure 4A, B) show that the pollen-dominance mechanism is retained in self-compatible Siberian *A. lyrata*. The same combination of S-alleles AhS1 (AkS-C) / AhS12 (AkS-D) in the F1 self-compatible accession (F1.1-1 plant in Table 2; Figure 4A) exists in *A. kamchatica* and is common in the eastern Siberian mountains bordering Okhotsk sea in Aldan-Amur interfluve (Figure 4C, yellow/blue pie charts). We, therefore, hypothesize that *A. kamchatica* with AhS1 (AkS-C) / AhS12 (AkS-D) combination of *S*-alleles was self-compatible in the first generation due to dominance of the AhS12 *S*-allele inherited from self-compatible Siberian *A. lyrata* over AhS1 inherited from *A. halleri*.

## Discussion

### Full *A. lyrata* genomes

Selfing accessions can be considered natural inbred lines which are especially useful in genomics, as the assembly of their genomes is not complicated by long heterozygous stretches. So far, only one selfing accession (MN47) of *A. lyrata* from North America has been fully assembled and serves as a reference for this species (Hu et al. 2011). An additional draft assembly of *A. lyrata* subsp. *petraea* has also been released (Paape et al. 2018), though its utility is hindered due to gaps in the assembly (12.75% missing) and lack of contiguity (scaffold N50 of 1.2Mb). Furthermore, while a single reference genome provides a useful resource for short-read re-sequencing-based population genetic studies (Novikova et al. 2016; The 1001 Genomes Consortium 2016), reference bias is an increasingly recognized problem. Using long and proximity-ligation reads we assembled high-quality genomes of the Siberian selfing *A. lyrata* accession NT1 and re-assembled North American *A. lyrata* MN47 accession. We found five inversions ranging from 0.3 to 2.4 Mb in length in between these independently evolved selfing accessions (Fig. 1, Supplementary Table 3). Large genomic structural rearrangements, especially inversions, can prevent chromosomal pairing and drive reproductive isolation and speciation (Rieseberg 2001; Stevison et al. 2011; McGaugh and Noor 2012; Ayala et al. 2013; Jeffares et al. 2017). In these circumstances selfing probably increases tolerance to such rearrangements and can even promote their fixation. For example, karyotypic changes from 8 to 5 chromosomes in *A. thaliana* are linked to a transition to self-compatibility at about 500 Kya (Durvasula et al. 2017). *A. lyrata* transitions to selfing are more recent but are consistent with this observation. Interestingly, the inversions found within *A. thaliana* (Jiao and Schneeberger 2020; Goel and Schneeberger 2022) and within *A. lyrata* (this study) are comparable in size: up to 2.5 Mb and 2.4 Mb respectively. However, to corroborate that selfing genomes are more tolerant to large structural rearrangements one must compare the results to outcrossing genomes which are not yet available at comparable quality, as heterozygosity renders them harder to assemble.

### Self-compatibility evolved at least twice in *A. lyrata*

We described the evolutionary history and distribution of selfing *A. lyrata* populations in Siberia, which have an independent origin from North American selfing *A. lyrata*. Siberian selfing populations posess only a single *S*-haplotype, AhS12 (AlS42), whereas several different haplogroups (AhS1 (AlS1), AhS31 (AlS19) and AhS29 (AlS13)) are found in the North American selfing populations of *A. lyrata* (Hu et al. 2011; Mable et al. 2017). The differences in *S*-haplotype composition of selfing lineages in Siberia and North America supports their independent origin, consistent with the phylogenetic relationships among accessions from these two regions (Supplementary Figure 13B). Our phylogenetic inference yields a well-supported clade of North American *A. lyrata*, comprised of both self-compatible and self-incompatible accessions, showing that the closest relatives to self-compatible North American *A. lyrata* are outcrossing North American *A. lyrata*, instead of self-compatible Siberian *A. lyrata*.

A transition to selfing is often associated with changes in flower morphology (Sicard and Lenhard 2011; Tsuchimatsu and Fujii 2022) which we observed in Siberian but not in North American selfing accessions (Supplementary Fig. 1C-E). The lack of so-called “selfing syndrome” in the latter was described previously (Carleial et al. 2017). Similarly, in the outcrossing species *Leavenworthia alabamica*, two independent selfing lineages have been described, with the older (∼150 Kya) showing an obvious selfing syndrome whereas the younger selfing lineage (∼48 Kya) did not (Busch et al. 2011). Although further investigation is required to quantify this difference in flower size, such observations in *A. lyrata* may also be explained by differences in transition to selfing: the North American *A. lyrata* likely transitioned to selfing during or after colonization of the area, around ∼10 Kya (Carleial et al. 2017), which is much more recent than our estimates of the Siberian selfer originating ∼90Kya.

### Transition to self-compatibility in Siberian *A. lyrata* is associated with S-locus

All selfing Siberian accessions spanning the massive geographical area between Lake Taymyr and Chukotka share the same *S*-haplotype (AhS12), suggesting the breakdown of self-incompatibility in Siberia is linked to the *S*-locus. This also suggests a single breakdown of self-incompatibility in the Siberian selfing lineage, as it is unlikely that this transition to self-compatibility occurred independently in multiple individuals with the same AhS12 allele. More than one origin of self-compatibility in the studied Siberian populations is improbable for two reasons: first, the *S*-locus is highly diverse, as tens of divergent *S*-alleles typically segregate within outcrossing populations to facilitate reproductive success (Schierup et al. 2001; Castric and Vekemans 2004; Castric and Vekemans 2007), so one would expect more diversity of *S*-alleles if there were multiple origins. And second, because dominant alleles, including AhS12, are more rare compared to recessive ones (Schierup et al. 1997; Billiard et al. 2007; Genete et al. 2020). The probability of independent loss of function on two of the same rare alleles is low (multiplied probabilities of drawing the same rare allele by chance).

However, the breakdown of self-incompatibility in North American *A. lyrata* is not associated with a specific *S*-allele or *S*-locus mutation, but rather with another genetic factor, likely in a downstream cascade of reactions preventing pollen-tube growth (Mable et al. 2005; Foxe et al. 2010; Mable et al. 2017). Therefore, despite strong evidence supporting an *S*-locus-driven loss of self-incompatibility in Siberian *A. lyrata*, it is possible that a mutation in a downstream cascade caused the initial mating system switch (Goring et al. 2014; Jany et al. 2019), followed by fixation of a single *S*-allele due to drift, and further degeneration of the *S*-allele sequence, reinforcing self-compatibility. Another scenario could involve a modifier mutation specific to AhS12 *S*-allele which arose prior to loss-of-function in the *S*-locus. The existence of allele-specific modifiers has been proposed based on observed segregation patterns in offspring (Nasrallah et al. 2004; Sherman-Broyles et al. 2007; Mable et al. 2017; Li et al. 2019) and could also explain loss of self-incompatibility in Siberian *A. lyrata* lineage fixed for the AhS12 *S*-allele. While these alternative explanations are plausible, based on the strong association between a specific S-haplotype (AhS12) and self-compatibility, we conclude that inactivation of AhS12 is the most likely scenario.

### Self-compatibility in Siberian *A. lyrata* is likely male-driven

Our long-read-based genome assembly of *A. lyrata* NT1 contains a fully-assembled *S*-locus (Figure 2), in which we manually annotated *SCR* by BLAST analysis of all known *SCR* sequences in *Arabidopsis*. The *SRK* gene was absent from our assembly. Mapping of the short reads from the *A. lyrata* NT1 accession to *A. halleri* AhS12 sequence of the same haplotype also did not yield any coverage of the *SRK* gene, so we conclude that *SRK* was lost from the NT1 genome. However, this does not mean that the loss of *SRK* is the causal mutation leading to selfing, as the SCR protein of NT1 *A. lyrata* also appears to be non-functional: (1) it lacks one of the eight cysteine residues (Figure 2C) that were shown to be functionally important (Kusaba et al. 2001; Mishima et al. 2003; Tsuchimatsu et al. 2010) (Figure 2C), and (2) its expression was not detected in flowers (Supplementary Fig. 9). Genotyping of the *S*-locus in other selfing *A. lyrata* accessions reveals that all of them share the same *S*-haplotype AhS12 (Figure 3C, Supplementary Table 1), which suggests their shared origin. Moreover, one of the selfing *A. lyrata* accessions has *SRK*, but seems to lack *SCR* (accession number MW0079456, Figure 3). Different reciprocal gene loss mutations of SCR or SRK across accessions (Figure 3B) exclude the possibility of gene loss being a causal mutation, and rather suggest that gene loss happened after a common causal mutation.

In controlled crossing experiments (Tsuchimatsu et al. 2012), haplogroup-D *SRK* in the *A. lyrata* subgenome of *A. kamchatica* (AkSRK-D, orthologous to AhSRK12) was shown to be functional. This suggests that *SRK* in the ancestors of both *A. kamchatica* and selfing *A. lyrata* was also functional. We discuss the role of selfing *A. lyrata* in the origin of *A. kamchatica* in the next section. If the breakdown of self-incompatibilty is indeed *S*-locus driven (and not caused by an unlinked *S*-allele specific modifier) it most likely occurred on *SCR* rather than *SRK* in this lineage. Our results show that indeed, SCR from NT1 is not recognised by a functional SRK of the same haplogroup (from accession NT8.4-24; Supplementary Fig. 11). Whether the initial loss-of-function in the SCR protein was due to a loss of a structurally important cysteine residue (Figure 2C) or a loss of expression (Supplementary Fig. 9A-B) is unclear. Transitioning to selfing through degradation of male specificity gene would be consistent with the recurrent pattern in the evolution of self-compatibility (reviewed in (Shimizu and Tsuchimatsu 2015). According to Bateman’s principle, an S-haplotype with non-functional SCR and functional SRK will produce pollen of higher fitness, as it will be compatible with all other S-haplotypes including itself. In contrast, an S-haplotype with a functional SCR and a non-functional SRK will produce pollen that will be self-compatible but incompatible with the fraction of the population carrying the same, albeit fully functional, S-haplotype.

Pistils with a non-functional SRK do not have a higher fitness unless pollen availability is very limited, making fixation of the male-driven selfing more likely (Bateman 1954; Tsuchimatsu and Shimizu 2013). The most likely scenario suggested by our results, where self-compatibility in Siberian *A. lyrata* is SCR-driven, is therefore consistent with Bateman’s principle.

### Self-compatible Siberian *A. lyrata* is ancestral to *A. kamchatica*

A previous study showed a Siberian *A. lyrata* accession (lyrpet4) to be genetically closest to *A. kamchatica*, however this was limited to sampling in a single locality and did not include assessment of *S*-alleles (Shimizu-Inatsugi et al. 2009; Paape et al. 2018). In addition to the previously reported selfing individual, our field and herbarium collections yielded seven more self-compatible accessions, spanning a wide geographical range across Siberia (Supplementary Table1). We explored the relationships among all Siberian *A. lyrata* accessions with *A. kamchatica* using network analysis and hierarchical clustering. Genetic network of Nei’s D (Figure 3B) shows that *A. kamchatica* clusters closely to self-compatible Siberian *A. lyrata*, which is consistent with the sister relationship between *A. kamchatica* and self-compatible *A. lyrata* in a well-supported maximum likelihood phylogeny (Supplementary Figure 13B). Moreover we identified a fixed S-allele (AhS12) associated with self-compatibility in Siberian *A. lyrata*.

Allopolyploid *A. kamchatica* has three *S*-alleles inherited from *A. halleri -* AhS26 (AkS-A), AhS47 (AkS-B) and AhS1 (AkS-C), and two *S*-alleles inherited from *A. lyrata* - AhS12 (AkS-D) and AhS02 (AkS-E) (Tsuchimatsu et al. 2012). The AhS12 *S*-allele is the most frequent in the *A. lyrata* subgenome of *A. kamchatica* and was inherited from a self-compatible Siberian *A. lyrata* lineage. A tree of *A. lyrata* and *A. kamchatica* accessions which share the AhS12 haplotype (based on exon 1 of the *SRK* gene; Supplementary Fig. 13A), shows a self-compatible *A. lyrata* accession is nested within a clade of *A. kamchatica* accessions, providing further support for their shared origin.

Furthermore, our demographic modeling suggests the Siberian selfing lineage originated approximately 90kya. This is in line with estimates by Paape et al. (2018), who dated the divergence times of both *A. kamchatica* subgenomes. Their estimates for divergence time of the *A. halleri* subgenome range from ∼60-100Kya, and the *A. lyrata* subgenome between ∼70-140Kya. The authors recommend caution when interpreting these parameters, and we agree: mutation rates used in both studies are from *A. thaliana* rather than *A. lyrata*, and sample sizes are small in both cases. Still, given the overlap in divergence estimates from both our study and work by Paape et al. (2018), it is plausible that at least one of the multiple polyploid origins of *A. kamchatica* included this selfing Siberian *A. lyrata* lineage as a parental genome donor.

Combinations of *A. kamchatica S*-alleles show a strong population structure (Figure 4C) consistent with multiple origins of *A. kamchatica* in different geographical regions (Shimizu et al. 2005; Shimizu-Inatsugi et al. 2009; Tsuchimatsu et al. 2012; Paape et al. 2018). However, the current sampling of *A. kamchatica* is biased towards Japan and the Kamchatka Peninsula, and this uneven coverage of the species range means observed frequencies of *S*-allele combinations may not represent their true distribution. That said, a combination of dominant non-functional AhS12 (*A. lyrata*-derived) and recessive AhS1 (*A. halleri*-derived) *S*-alleles is common in *A. kamchatica* in the eastern Siberian mountains bordering Okhotsk sea in Aldan-Amur interfluve (Figure 4C).

Interestingly, while both progenitors of *A. kamchatica* co-exist in Europe, and interspecific crosses can be created *ex situ* (Sarret et al. 2009), *A. lyrata* and *A. halleri* do not form other allotetraploids (Clauss and Koch 2006; Schmickl et al. 2010)). The variation (or lack thereof) of mating systems in *A. lyrata* and *A. halleri* can explain why allopolyploid establishment is limited to Asia: *A. halleri* is self-incompatible throughout its range (no known selfing accessions have been described to date), and selfing *A. lyrata* is found only in Siberia and North America. Previous work showed that self-compatibility in *A. kamchatica* was likely male (SCR)-driven in the more dominant S-haplotype inherited from *A. lyrata* (Ah12/Al42/Ak-D) (Tsuchimatsu et al. 2012). We argue that self-compatibilty is ancestral to *A. kamchatica*, and inherited from Siberian *A. lyrata*. We also show that dominance between non-functional AhS12 and functional AhS01 is retained in self-compatible *A. lyrata* (Figure 4A,B) and therefore argue the transition to selfing in *A. kamchatica* with this combination of *S*-alleles was likely immediate upon allopolyploid formation. Our results show that Siberian selfing diploid *A. lyrata* is ancestral to allotetraploid *A. kamchatica*, and contribued the most widely observed *A. lyrata*-derived *S*-allele (AhS12) in *A. kamchatica*. Furthermore, the non-functional AhS12 *S*-allele is still dominant over the recessive AhS01 *S*-allele in *A. lyrata.* This dominance of the non-functional *S*-allele likely explains the transition to self-compatibility in *A. kamchatica* with the same combination of *S*-alleles (AhS12/AkS-D and AhS01/AkS-C), rather than self-compatibility evolving *de novo* in *A. kamchatica*.

Similar examples where a loss-of-function mutation on a dominant S-haplotype in one progenitor facilitated transition to selfing in allotetraploids have been recently reviewed (Novikova et al. 2022) and include *A. suecica* (Novikova et al. 2017), *Capsella bursa-pastoris* (Bachmann et al. 2019; Bachmann et al. 2021; Duan et al. 2023) and *Brassica napus* (Okamoto et al. 2007; Kitashiba and Nasrallah 2014). Allopolyploid establishment may be facilitated by a transition to self-compatibility, ensuring reproductive success in the face of limited mating partners.

## Materials and Methods

### Plant collection and growth

We collected seeds from three *A. lyrata* populations (NT1, NT8 and NT12) during an expedition to the Yakutia region in Russia in the summer of 2019 (Supplementary Table 1). Multiple individual plants were collected from those three populations: three individuals from NT1 (NT1_1, NT1_2, NT1_3), four from NT8 (NT8_1, NT8_2, NT8_3, NT8_4) and two from NT12 (NT12_1, NT12_2). Collected seeds were grown in the greenhouse at 21°C, under 16 hours of light per day until a full rosette was formed, after which plants were moved to open frames outside on the grounds of the Max Planck Institute for Plant Breeding Research in Cologne, Germany. We grew several seeds per collected bag of seeds from individual plants, each was given an additional number extension (e.g., NT1_1**_1,** NT1_1**_2**, etc.). In this work we only used the plants with last extension 1. All the individuals grown from NT1 population formed long fruits and appeared to be selfing. NT1 samples were collected on a sandy island in the course of the Lena river (GPS coordinates 66.80449, 123.46546; Supplementary Figure 1A shows a picture of the collection site).

### Pollen tube staining to characterize mating type

Almost mature flower buds were opened and, after removing the anthers, manually pollinated. Pistils were collected 2–3 hours after pollination, fixed for 1.5 hours in 10% acetic acid in ethanol and softened in 1 M NaOH overnight. Before staining, the tissue was washed three times in KPO_4_ buffer (pH 7.5). For staining we submerged the tissue in 0.01% aniline blue for 10–20 minutes. After that, pistils were transferred to slides into mounting media and observed under UV light (Lu 2011). A self-compatible reaction was called if we counted more than 10 pollen tubes.

#### Long-read sequencing for *de novo* genome assembly

DNA extraction, library preparation and long-read sequencing of the NT1 and MN47 *A. lyrata* accessions were performed by the Max Planck-Genome-centre Cologne, Germany https://mpgc.mpipz.mpg.de/home/). High molecular weight DNA was isolated from 1.5 g material with a NucleoBond HMW DNA kit (Macherey Nagel). Quality was assessed with a FEMTOpulse device (Agilent) and quantity measured by a Quantus fluorometer (Promega). HiFi libraries were then prepared according to the manual “Procedure & Checklist - Preparing HiFi SMRTbell® Libraries using SMRTbell Express Template Prep Kit 2.0” with an initial DNA fragmentation by g-Tubes (Covaris) and final library size selection on BluePippin (Sage Science). Size distribution was again controlled by FEMTOpulse (Agilent). Size-selected libraries were then sequenced on a Sequel II device with Binding Kit 2.0 and Sequel II Sequencing Kit 2.0 for 30 h (Pacific Biosciences).

#### Short-read sequencing for population analyses

Plant material was processed in two different ways, indicated by type I and II in Supplementary Table 1. Type I: herbarium material was extracted in a dedicated clean-room facility (Ancient DNA Laboratory, Department of Archaeology, University of Cambridge). The lab has strict entry and surface decontamination protocols, and no nucleic acids are amplified in the lab. For each accession, leaf and/or stem tissue was placed in a 2 mL tube with 2 tungsten carbide beads and ground to a fine powder using a Qiagen Tissue Lyser. Each batch of extractions included a negative extraction control (identical but without tissue). DNA was extracted using the DNeasy Plant Mini Kit (Qiagen). Libraries preparation and sequencing were performed by Novogene LTD (UK). Sequencing libraries were generated using NEBNext® DNA Library Prep Kit following manufacturer’s recommendations and indices were added to each sample. The genomic DNA is randomly fragmented to a size of 350bp by shearing, then DNA fragments were end polished, A-tailed, and ligated with the NEBNext adapter for Illumina sequencing, and further PCR enriched by P5 and indexed P7 oligos. The PCR products were purified (AMPure XP system) and resulting libraries were analyzed for size distribution by Agilent 2100 Bioanalyzer and quantified using real-time PCR.

Type II: Genomic DNA was isolated with the “NucleoMag© Plant ’’ kit from Macherey and Nagel (Düren, Germany) on the KingFisher 96Plex device (Thermo) with programs provided by Macherey and Nagel. Random samples were selected for a quality control to ensure intact DNA as a starting point for library preparation. TPase-based libraries were prepared as outlined by (Rowan et al. 2019) on a Sciclone (PerkinElmer) robotic device. Short-read (PE 150bp) sequencing was performed by Novogene LTD (UK), using a NovaSeq 6000 S4 flow cell Illumina system.

#### Transcriptome sequencing for *S*-locus gene expression assessment

We used three flash-frozen open flowers of the *A. lyrata* NT1 accession as input material for RNA sequencing, which we used to assess the expression of the *S*-locus genes. RNA was extracted by the RNeasy Plant Kit (Qiagen) including an on-column DNase I treatment. Quality was assessed by Agilent Bioanalyser and the amount was calculated by an RNA-specific kit for Quantus (Promega). An Illumina-compatible library was prepared with the NEBNext® Ultra™ II RNA Library Prep Kit for Illumina ® and finally sequenced on a HiSeq 3000 at the Max Planck-Genome-centre Cologne.

#### PacBio *de novo* assembly and annotation of NT1 and MN47 *A. lyrata* accessions

Raw PacBio reads of NT1 were assembled using Hifiasm assembler (Cheng et al. 2021) in the default mode, choosing the primary contig graph as our resulting assembly. The completeness of our assembly was assessed using BUSCO (Seppey et al. 2019) with Brassicales_odb10 set. Repeated sequences were masked using RepeatMasker (Smit et al. 2013-2015) with the merged libraries of RepBase *A. thaliana* repeats and NT1 *A. lyrata* repeats, which we modeled with RepeatModeler (Smit and Hubley 2008-2015). Then, annotation from the reference MN47 genome (Rawat et al. 2015) was transferred to our NT1 repeat-masked assembly by using Liftoff (Shumate and Salzberg 2020). Contigs were reordered according to their alignment to the reference chromosomes and updated gene and repeat annotations using RagTag (Alonge et al. 2019) in the scaffolding mode without correction. Assembly of MN47 PacBio reads was done using the Hifiasm assembler with the same parameters.

#### Synteny analysis of *A. lyrata*, *A. suecica* and *C. rubella* genomes

Synteny analysis was done by performing an all-against-all BLASTp search using the CDS sequences of both genomes. We used SynMap (Haug-Baltzell et al. 2017), a tool from the online platform CoGe, with the default parameters for DAGChainer. The Quota Align algorithm was used to decide on the syntenic depth, employing the default parameters. Syntenic blocks were not merged. The results were visualized using the R (version 4.1.2) library ‘circlize’ (version 0.4.13), as well as using plotsr (version 0.5.3) (Goel and Schneeberger 2022) for the supplementary figures.

#### HiC sequencing of NT1 *A. lyrata* accession to validate structural variants

A chromatin-capture library of the NT1 *A. lyrata* accession was prepared by the Max Planck-Genome-centre Cologne, Germany, and was used for validation of the large inversions in whole-genome comparisons. We followed the Dovetail® Omni-C® Kit starting with 0.5 g of fresh weight as input. Libraries were quantified and quality assessed by capillary electrophoresis (Agilent Tapestation) and then sequenced at the Novogene Ltd (UK), using a NovaSeq Illumina system.

#### Mapping of Hi-C reads for the *A. lyrata* accessions NT1 and MN47

To validate the assembled scaffolds of *A. lyrata*, we used proximity-ligation short read Hi-C data. For NT1, Hi-C reads were mapped to the repeat-masked NT1 genome assembly, using the mapping pipeline proposed by the manufacturer (https://omni-c.readthedocs.io/en/latest/index.html). The Dovetail Omni-C processing pipeline is based on BWA (Li and Durbin 2009), pairtools (https://github.com/mirnylab/pairtools) and Juicertools (Durand et al. 2016). We mapped the Hi-C reads for MN47 (released previously (Zhu et al. 2017)) to a repeat masked MN47 genome (Hu et al. 2011) and to a repeat masked version of the newly assembled MN47 genome (in this paper) using HiCUP (version 0.6.1) (Wingett et al. 2015). The assemblies were manually examined using Juicebox (Robinson et al. 2018). Plots of the HiC contact matrix were made using the function hicPlotMatrix from HiCExplorer (Wolff et al. 2020) (version 3.7.2).

#### Validation of structural variants between NT1 and MN47 *A. lyrata* accessions

To validate the inversions (Supplementary Table 2) we used PacBio, Hi-C data and synteny analysis results. Guided by synteny analyses, we first identified inversion breakpoints. Then, we investigated the long read map at these regions and either confirmed their contiguity or manually flipped the genomic region, followed by another round or long read map investigation (Supplementary Fig 3-8). To map the PacBio HiFi reads we used Winnowmap (Jain et al. 2020). As the last step, we analyzed the Hi-C contact maps in the same regions to show that there is no evidence for alternative genome assembly configurations (Supplementary Figure 3-7).

#### *A. lyrata* NT1 *S*-locus genotyping and manual annotation

We manually annotated the *S*-locus in our initial assembly before the reference-guided reordering and scaffolding. In the transferred annotation resulting from Liftoff (Shumate and Salzberg 2020) we found both of the flanking genes (*U-box* and *ARK3*) in the same contig. The final coordinates of the *S*-locus in the NT1 assembly on scaffold 7 are 9,291,658bp to 9,336,246bp. The length of the assembled NT1 *A. lyrata S*-locus including both flanking genes is about 44.5Kbp. We mapped PacBio long reads back to the assembled NT1 genome using minimap2 (Li 2018) with default parameters in order to make sure that there are no obvious gaps in coverage or break points (Supplementary Fig. 8). Similarly to Zhang et al. (Zhang et al. 2019), we blasted the *SRK* and *SCR* sequences from all the known *S*-haplotypes across *Arabidopsis* and *Capsella* to the *A. lyrata* NT1 *S*-locus, finding a single hit at the *SCR* gene from the AhS12 haplogroup. We constructed a comparative structure plot of *A. lyrata* NT1 and *A. halleri* S12 (Genbank accession KJ772374) *S*-loci (Fig. 2B) using the R library genoPlotR (Guy et al. 2010). We aligned SCR protein sequences using MAFFT with default parameters and estimated a phylogenetic tree with RaxML (Stamatakis 2014) using the BLOSUM62 substitution model and visualized the alignment (Fig. 2C) using Jalview2 (Waterhouse et al. 2009). The phylogenetic tree was visualized using R package “ape” (Paradis et al. 2004).

#### *A. lyrata S*-allele genotyping from short-read sequencing data

The *S*-alleles from all the re-sequenced samples used in the population analysis (Supplementary Table 1) and crosses (NT8.4-24, submitted to ENA under ERS12276051) were genotyped using the *S*-locus genotyping pipeline NGSgenotyp (Genete et al. 2020). The list of *SRK* and *SCR* alleles used as a reference dataset is provided in the Supplementary Table 3 and the corresponding sequences for *SRK* and *SCR* alleles are provided in the Supplementary Data 1 and 2. Using the NGSgenotyp pipeline we could not identify any S-haplotypes for DRR124344 (lyrpet4), for either *SRK* or *SCR* databases. However, we found a partial *SCR* gene sequence matching the AhS12 haplotype by blasting the *SCR* database to the DRR124344 assembly. We translated the SCR nucleotide sequence and aligned the resulting protein sequence with SCR proteins from other accessions using MAFFT (Katoh and Standley 2013) using default parameters. The resulting alignment shows that SCR from DRR124344 is shorter compared to NT1 or AhS12. To confirm that SCR from DRR124344 belongs to the AhS12 haplotype, we estimated a maximum likelihood tree using IQ-tree web service (http://www.iqtree.org/) with default parameters (Supplementary Fig. 10B).

#### Short read mapping and variant calling for population analysis

We first filtered the short paired-end reads (2×150bp) for adapter contamination using bbduk.sh script from BBMap (38.20) (Bushnell 2014) with the following parameters settings: ktrim=r k=23 mink=11 hdist=1 tbo tpe qtrim=rl trimq=15 minlen=70. Then we mapped the reads to the MN47 and NT1 *A. lyrata* genome with bwa mem (0.7.17) (Li and Durbin 2009), marking shorter split reads as secondary (-M parameter). We marked potentially PCR duplicated reads with picard MarkDuplicates (http://broadinstitute.github.io/picard/), sorted and indexed the bam file with samtools (Li et al. 2009). To call variants, we used the HaplotypeCaller algorithm from GATK (McKenna et al. 2010) (3.8). We then ran GenotypeGVCF from GATK including non-variant sites on the entire sample set to generate a vcf. To estimate heterozygosity levels, we calculated the proportion of heterozygous sites within all the confidently called sites in mapping to MN47 and NT1 reference genomes (Supplementary Table 1, Supplementary Fig. 12).

#### Separation of subgenomes from *A. kamchatica* accessions

To isolate the A. *lyrata* subgenome of *A. kamchatica*, we used a combined reference, containing *A. lyrata* NT1 and *A. halleri ssp. gemmifera* reference genomes (Briskine et al. 2016). We mapped *A. kamchatica* short reads to the combined reference with bwa mem (0.7.17) (Li and Durbin 2009) and filtered for reads mapped uniquely to *A. lyrata* NT1 using samtools (Li et al. 2009). We then genotyped the resulting *A. lyrata*-subgenome bam files for each *A. kamchatica* accession as described above for diploid samples.

#### Tree and network estimation

##### Genome-wide SNP tree

We filtered the vcf generated above to include only biallelic SNPs without missing data, which resulted in 2,261,679 SNPs. These data were read into R (version 4.1.1) and from them we estimated a neighbor-joining tree using the nj function from package ape (Paradis and Schliep 2019). We then visualized the neighbor-joining tree as a cladogram using ggtree (Yu et al. 2017; Yu et al. 2018; Yu 2020) and annotated the tips with associated data (Supplementary Figure 13B). We then further filtered this dataset to include only Siberian *A. lyrata* and an outgroup (excluding *A. kamchatica* from this portion) to generate the *lyrata*-only tree (Figure 3C).

##### Network based on Nei’s D and phylogenetic inference

We filtered a vcf of biallelic SNPs shared by the *lyrata* subgenome of *A. kamchatica* and all *A. lyrata* accessions down to just four-fold degenerate sites, with maximum 10% missing data across individuals, resulting in 4,141 SNPs. We read the vcf with both Siberian *A. lyrata* and *A. kamchatica* into R using vcfR (Knaus and Grünwald 2017), then calculated Nei’s D (Nei 1972) between individuals using StAMPP (Pembleton et al. 2013). We visualized the resulting matrix in SplitsTree4 (Huson and Bryant 2006) and in R using the pheatmap package (Kolde 2019). To further explore the evolutionary relationships among accessions, we generated a nexus file from the vcf using vcf2phylip (Ortiz 2019) which served as input for phylogenetic inference with IQTree (http://www.iqtree.org/)

##### SRK tree

We assembled partial *SRK* sequences from Siberian *A. lyrata* and *A. kamchatica* accessions based on short-read sequencing data using the assembly step of the *S*-locus genotyping pipeline NGSgenotyp (Genete et al. 2020) and aligned sequences with MAFFT (Katoh and Standley 2013). From this alignment estimated 1000 bootstrap replicates of a maximum likelihood phylogeny using RaXML (Stamatakis 2014) with substitution model GTR+**Γ,** then visualized the best-scoring ML phylogeny using R package ape 5.0 (Paradis and Schliep 2019). The input alignment is available in Supplementary Data 3.

##### PCR identification of AhS12 haplotype

For DNA extraction, 1cm of leaf material was frozen in liquid nitrogen and ground to a powder. We added 400 µl UltraFastPrep Buffer to the powdered tissue, then mixed, vortexed, and finally spun for five minutes at 5000 rpm. We then took 300 µl of the supernatant, added 300 µl isopropanol, and mixed by inversion. We again spun for five minutes at 5000 rpm, then discarded the supernatant and dried 10-30 minutes at 37°C. The pellet was resuspended in 200 µl 1xTE and stored at 4°C. We amplified the AhSRK12 allele by PCR using 1.5 µl of DNA solution and previously published primers (forward ATCATGGCAGTGGAACACAG, reverse CAAATCAGACAACCCGACCC) (Ruggiero et al. 2008). We ran 35 cycles consisting of 30 s at 94°C, 30 s annealing at 56.8°C, and a 40 s extension at 72°C. We visualized PCR products via gel electrophoresis using 1.5% agarose gel with GelGreen® nucleic acid stain (Supplementary figure 10A). Accessions identified with SRK 12 (NT8.4-24) and without SRK 12 (NT8.4-20) were used in crosses (Supplementary Figure 10B-D).

#### Demographic modeling of divergence between selfing and outcrossing Siberian *A. lyrata* lineages

We calculated nucleotide diversity using all biallelic and non-variant sites in 10kb windows with custom script uploaded to github (https://github.com/novikovalab/selfing_Alyrata) Confidence intervals for the median of the distribution were calculated using the basic bootstrap method in the R package ‘boot’(Davison and Hinkley 1997; Canty and Ripley 2022).

To prepare a joint allele frequency spectrum of the seven self-compatible accessions and the ten self-incompatible accessions, we first filtered the SNP-only vcf to remove centromeric, pericentromeric, and exonic regions. We subsequently filtered out sites with missing data to yield our final vcf for demographic inference. Following Nordborg & Donnelly (Nordborg and Donnelly 1997) we excluded sites heterozygous in the selfing population and treated selfers as haploid. We then generated the joint allele frequency spectrum using easySFS (https://github.com/isaacovercast/easySFS). EasySFS produces output ready for use in fastsimcoal2 (fsc26) (Excoffier et al. 2013; Excoffier et al. 2021), which we then used for demographic modeling. We tested five models for the origin of self-compatibility in Siberian *A. lyrata*, (A) simple divergence, (B) divergence with symmetrical introgression (migration) (C) divergence with asymmetrical introgression, (D) Simple divergence model as in Model A, plus bottleneck in selfing population. (E) Model C (asymmetric gene flow) plus bottleneck in selfing population.

For each model, we initiated 100 fastsimcoal2 runs. We then chose the best run for each model (the run with the best likelihood scores) and from that best run calculated the Aikake Information Criterion for the model. After selecting the model with the best AIC score, we used the maximum likelihood parameter file to generate 200 pseudo observations of joint SFS for bootstrapping. For each of the 200 pseudo observations, we initiated 100 fastsimcoal2 runs, then selected the best run for each model based on likelihood scores as above. The resulting parameter estimates from the 200 replicate pseudo-observations were used to calculate the 95% confidence intervals in R. Site frequency spectra and other fastsimcoal2 input files (.tpl and .est) are on GitHub (https://github.com/novikovalab/selfing_Alyrata). Because fastsimcoal2 reports haploid effective population sizes, we divided them by two to report numbers of diploid individuals (Table 1). These parameters can be interpreted as the inverse of the coalescent rate estimated from our accessions.

## Acknowledgements

We thank the Max Planck-Genome-centre Cologne (http://mpgc.mpipz.mpg.de/home/) for performing short- and long-read libraries and PacBio sequencing in this study. We thank Vincent Castric and Mathieu Genete for sharing the up-to-date *SRK* database for the *S*-locus genotyping pipeline. We thank Kathrin Wippel for providing seeds of North American MN47 *A. lyrata*. We thank Jo Osborn, University of Cambridge, for assisting with the herbarium extractions. The work of A.P. Seregin on curation of dry plant material was supported by the Russian Science Foundation (project #21-77-20042). Field research was supported by the Austrian Science Fund (FWF grant P30208-B-29) and held within the state assignment of the Papanin Institute for Biology of Inland Waters Russian Academy of Sciences (theme 121051100099-5). The sequencing and data analysis was funded by the European Union (ERC, HOW2DOUBLE, 101041354) and by the Deutsche Forschungsgemeinschaft (DFG) - Project number 462181533. Views and opinions expressed are however those of the author(s) only and do not necessarily reflect those of the European Union or the European Research Council Executive Agency. Neither the European Union nor the granting authority can be held responsible for them.

## Data availability

The whole genome raw Illumina short reads for the samples used in this study were submitted to the ENA database under the project number PRJEB50329 (ERP134897). Individual accession names are listed in the Supplementary Table 1. Raw PacBio HiFi reads of NT1 and MN47, Hi-C reads of NT1, RNAseq reads of NT1, and the genome assembly and annotation of *A. lyrata* NT1 (GCA_945152055) and MN47 (GCA_944990045) have been submitted to ENA database under the same project number PRJEB50329 (ERP134897) and to https://figshare.com/projects/Arabidopsis_lyrata_genome_assemblies/162343 Scripts associated with the project are at https://github.com/novikovalab/selfing_Alyrata.

## Author contributions

UKK, ADS, and PYN conceptualized the project. UKK, NPT, UP, ACC, APS, and LY collected material and/or generated data. UKK, ADS, JDVdV, RB, NPT, XV, SL, and PYN analyzed the data. UKK, ADS, JDVdV, RB, and PYN wrote the manuscript. All authors read and approved the final manuscript.

## References

1. Alonge M, Soyk S, Ramakrishnan S, Wang X, Goodwin S, Sedlazeck FJ, Lippman ZB, Schatz MC. 2019. RaGOO: fast and accurate reference-guided scaffolding of draft genomes. Genome Biol. 20:224.

2. Ayala D, Guerrero RF, Kirkpatrick M. 2013. Reproductive isolation and local adaptation quantified for a chromosome inversion in a malaria mosquito. Evolution 67:946–958.

3. Bachmann JA, Tedder A, Fracassetti M, Steige KA, Lafon-Placette C, Köhler C, Slotte T. 2021. On the origin of the widespread self-compatible allotetraploid Capsella bursa-pastoris (Brassicaceae). Heredity [Internet]. Available from: http://dx.doi.org/10.1038/s41437-021-00434-9

4. Bachmann JA, Tedder A, Laenen B, Fracassetti M, Désamoré A, Lafon-Placette C, Steige KA, Callot C, Marande W, Neuffer B, et al. 2019. Genetic basis and timing of a major mating system shift in Capsella. New Phytol. 224:505–517.

5. Bateman AJ. 1954. Self-incompatibility systems in angiosperms II. Iberis amara. Heredity 8:305–332.

6. Bechsgaard J, Bataillon T, Schierup MH. 2004. Uneven segregation of sporophytic self-incompatibility alleles in Arabidopsis lyrata. J. Evol. Biol. 17:554–561.

7. Billiard S, Castric V, Vekemans X. 2007. A general model to explore complex dominance patterns in plant sporophytic self-incompatibility systems. Genetics 175:1351–1369.

8. Boggs NA, Dwyer KG, Shah P, McCulloch AA, Bechsgaard J, Schierup MH, Nasrallah ME, Nasrallah JB. 2009. Expression of distinct self-incompatibility specificities in Arabidopsis thaliana. Genetics 182:1313–1321.

9. Boggs NA, Nasrallah JB, Nasrallah ME. 2009. Independent S-locus mutations caused self-fertility in Arabidopsis thaliana. PLoS Genet. 5:e1000426.

10. Briskine RV, Paape T, Shimizu-Inatsugi R, Nishiyama T, Akama S, Sese J, Shimizu KK. 2016. Genome assembly and annotation of Arabidopsis halleri, a model for heavy metal hyperaccumulation and evolutionary ecology. Mol. Ecol. Resour. [Internet]. Available from: https://www.ncbi.nlm.nih.gov/pubmed/27671113

11. Burghgraeve N, Simon S, Barral S, Fobis-Loisy I, Holl A-C, Ponitzki C, Schmitt E, Vekemans X, Castric V. 2020. Base-Pairing Requirements for Small RNA-Mediated Gene Silencing of Recessive Self-Incompatibility Alleles in Arabidopsis halleri. Genetics 215:653–664.

12. Burns R, Mandáková T, Gunis J, Soto-Jiménez LM, Liu C, Lysak MA, Novikova PY, Nordborg M. 2021. Gradual evolution of allopolyploidy in Arabidopsis suecica. Nat Ecol Evol [Internet]. Available from: http://dx.doi.org/10.1038/s41559-021-01525-w

13. Busch JW, Joly S, Schoen DJ. 2011. Demographic signatures accompanying the evolution of selfing in Leavenworthia alabamica. Mol. Biol. Evol. 28:1717–1729.

14. Bushnell B. 2014. BBMap: A Fast, Accurate, Splice-Aware Aligner. Lawrence Berkeley National Lab. (LBNL), Berkeley, CA (United States) Available from: https://www.osti.gov/servlets/purl/1241166

15. Canty A, Ripley BD. 2022. boot: Bootstrap R (S-Plus) Functions.

16. Carleial S, van Kleunen M, Stift M. 2017. Small reductions in corolla size and pollen: ovule ratio, but no changes in flower shape in selfing populations of the North American Arabidopsis lyrata. Oecologia 183:401–413.

17. Castric V, Bechsgaard J, Schierup MH, Vekemans X. 2008. Repeated adaptive introgression at a gene under multiallelic balancing selection. Bergelson J, editor. PLoS Genet. 4:e1000168.

18. Castric V, Bechsgaard JS, Grenier S, Noureddine R, Schierup MH, Vekemans X. 2010. Molecular evolution within and between self-incompatibility specificities. Mol. Biol. Evol. 27:11–20.

19. Castric V, Vekemans X. 2004. Plant self-incompatibility in natural populations: a critical assessment of recent theoretical and empirical advances. Mol. Ecol. 13:2873–2889.

20. Castric V, Vekemans X. 2007. Evolution under strong balancing selection: how many codons determine specificity at the female self-incompatibility gene SRK in Brassicaceae? BMC Evol. Biol. 7:132.

21. Charlesworth D, Vekemans X, Castric V, Glémin S. 2005. Plant self-incompatibility systems: a molecular evolutionary perspective. New Phytol. 168:61–69.

22. Cheng H, Concepcion GT, Feng X, Zhang H, Li H. 2021. Haplotype-resolved de novo assembly using phased assembly graphs with hifiasm. Nat. Methods 18:170–175.

23. Clauss MJ, Koch MA. 2006. Poorly known relatives of Arabidopsis thaliana. Trends Plant Sci. 11:449– 459.

24. Davison AC, Hinkley DV. 1997. Bootstrap Methods and Their Applications. Available from: http://statwww.epfl.ch/davison/BMA/

25. Duan T, Zhang Z, Genete M, Poux C, Sicard A, Lascoux M, Castric V, Vekemans X. 2023. Dominance between self-incompatibility alleles determines the mating system of Capsella allopolyploids. bioRxiv [Internet]:2023.04.17.537155. Available from: https://www.biorxiv.org/content/10.1101/2023.04.17.537155v1

26. Dukić M, Bomblies K. 2022. Male and female recombination landscapes of diploid Arabidopsis arenosa. Genetics [Internet]. Available from: http://dx.doi.org/10.1093/genetics/iyab236

27. Durand E, Méheust R, Soucaze M, Goubet PM, Gallina S, Poux C, Fobis-Loisy I, Guillon E, Gaude T, Sarazin A, et al. 2014. Dominance hierarchy arising from the evolution of a complex small RNA regulatory network. Science 346:1200–1205.

28. Durand NC, Shamim MS, Machol I, Rao SSP, Huntley MH, Lander ES, Aiden EL. 2016. Juicer Provides a One-Click System for Analyzing Loop-Resolution Hi-C Experiments. Cell Syst 3:95–98.

29. Durvasula A, Fulgione A, Gutaker RM, Alacakaptan SI, Flood PJ, Neto C, Tsuchimatsu T, Burbano HA, Picó FX, Alonso-Blanco C, et al. 2017. African genomes illuminate the early history and transition to selfing in Arabidopsis thaliana. Proc. Natl. Acad. Sci. U. S. A. 114:5213–5218.

30. Dwyer KG, Berger MT, Ahmed R, Hritzo MK, McCulloch AA, Price MJ, Serniak NJ, Walsh LT, Nasrallah JB, Nasrallah ME. 2013. Molecular characterization and evolution of self-incompatibility genes in Arabidopsis thaliana: the case of the Sc haplotype. Genetics 193:985–994.

31. Excoffier L, Dupanloup I, Huerta-Sánchez E, Sousa VC, Foll M. 2013. Robust demographic inference from genomic and SNP data. PLoS Genet. 9:e1003905.

32. Excoffier L, Marchi N, Marques DA, Matthey-Doret R, Gouy A, Sousa VC. 2021. *fastsimcoal2*: demographic inference under complex evolutionary scenarios. Bioinformatics [Internet] 37:4882– 4885. Available from: http://dx.doi.org/10.1093/bioinformatics/btab468

33. Foxe JP, Stift M, Tedder A, Haudry A, Wright SI, Mable BK. 2010. Reconstructing origins of loss of self-incompatibility and selfing in North American Arabidopsis lyrata: a population genetic context. Evolution 64:3495–3510.

34. Fujii S, Takayama S. 2018. Multilayered dominance hierarchy in plant self-incompatibility. Plant Reprod. 31:15–19.

35. Fulgione A, Koornneef M, Roux F, Hermisson J, Hancock AM. 2018. Madeiran Arabidopsis thaliana Reveals Ancient Long-Range Colonization and Clarifies Demography in Eurasia. Mol. Biol. Evol. 35:564–574.

36. Genete M, Castric V, Vekemans X. 2020. Genotyping and De Novo Discovery of Allelic Variants at the Brassicaceae Self-Incompatibility Locus from Short-Read Sequencing Data. Mol. Biol. Evol. 37:1193–1201.

37. Goel M, Schneeberger K. 2022. plotsr: Visualising structural similarities and rearrangements between multiple genomes. bioRxiv [Internet]:2022.01.24.477489. Available from: https://www.biorxiv.org/content/10.1101/2022.01.24.477489v1

38. Goring DR, Indriolo E, Samuel MA. 2014. The ARC1 E3 ligase promotes a strong and stable self-incompatibility response in Arabidopsis species: response to the Nasrallah and Nasrallah commentary. Plant Cell 26:3842–3846.

39. Goubet PM, Berges H, Bellec A, Prat E, Helmstetter N, Mangenot S, Gallina S, Holl AC, Fobis-Loisy I, Vekemans X, et al. 2012. Contrasted patterns of molecular evolution in dominant and recessive self-incompatibility haplotypes in Arabidopsis. PLoS Genet. 8:e1002495.

40. Griffin PC, Willi Y. 2014. Evolutionary shifts to self-fertilisation restricted to geographic range margins in North American Arabidopsis lyrata. Ecol. Lett. 17:484–490.

41. Guo Y-L, Bechsgaard JS, Slotte T, Neuffer B, Lascoux M, Weigel D, Schierup MH. 2009. Recent speciation of Capsella rubella from Capsella grandiflora, associated with loss of self-incompatibility and an extreme bottleneck. Proc. Natl. Acad. Sci. U. S. A. 106:5246–5251.

42. Guo YL, Zhao X, Lanz C, Weigel D. 2011. Evolution of the S-locus region in Arabidopsis relatives. Plant Physiol. 157:937–946.

43. Guy L, Kultima JR, Andersson SG. 2010. genoPlotR: comparative gene and genome visualization in R. Bioinformatics 26:2334–2335.

44. Hatakeyama K, Takasaki T, Suzuki G, Nishio T, Watanabe M, Isogai A, Hinata K. 2001. The S receptor kinase gene determines dominance relationships in stigma expression of self-incompatibility in Brassica. Plant J. 26:69–76.

45. Haug-Baltzell A, Stephens SA, Davey S, Scheidegger CE, Lyons E. 2017. SynMap2 and SynMap3D: web-based whole-genome synteny browsers. Bioinformatics 33:2197–2198.

46. Henry IM, Dilkes BP, Tyagi A, Gao J, Christensen B, Comai L. 2014. The BOY NAMED SUE quantitative trait locus confers increased meiotic stability to an adapted natural allopolyploid of Arabidopsis. Plant Cell 26:181–194.

47. Huson DH, Bryant D. 2006. Application of phylogenetic networks in evolutionary studies. Mol. Biol. Evol. 23:254–267.

48. Hu TT, Pattyn P, Bakker EG, Cao J, Cheng J-F, Clark RM, Fahlgren N, Fawcett JA, Grimwood J, Gundlach H, et al. 2011. The Arabidopsis lyrata genome sequence and the basis of rapid genome size change. Nat. Genet. 43:476–481.

49. Jain C, Rhie A, Zhang H, Chu C, Walenz BP, Koren S, Phillippy AM. 2020. Weighted minimizer sampling improves long read mapping. Bioinformatics 36:i111–i118.

50. Jany E, Nelles H, Goring DR. 2019. The Molecular and Cellular Regulation of Brassicaceae Self-Incompatibility and Self-Pollen Rejection. Int. Rev. Cell Mol. Biol. 343:1–35.

51. Jeffares DC, Jolly C, Hoti M, Speed D, Shaw L, Rallis C, Balloux F, Dessimoz C, Bähler J, Sedlazeck FJ. 2017. Transient structural variations have strong effects on quantitative traits and reproductive isolation in fission yeast. Nat. Commun. 8:14061.

52. Jiao W-B, Schneeberger K. 2020. Chromosome-level assemblies of multiple Arabidopsis genomes reveal hotspots of rearrangements with altered evolutionary dynamics. Nat. Commun. 11:989.

53. Kamau E, Charlesworth D. 2005. Balancing selection and low recombination affect diversity near the self-incompatibility loci of the plant Arabidopsis lyrata. Curr. Biol. 15:1773–1778.

54. Katoh K, Standley DM. 2013. MAFFT multiple sequence alignment software version 7: improvements in performance and usability. Mol. Biol. Evol. 30:772–780.

55. Kitashiba H, Nasrallah JB. 2014. Self-incompatibility in Brassicaceae crops: lessons for interspecific incompatibility. Breed. Sci. 64:23–37.

56. Knaus BJ, Grünwald NJ. 2017. vcfr: a package to manipulate and visualize variant call format data in R. Mol. Ecol. Resour. 17:44–53.

57. Kodera C, Just J, Da Rocha M, Larrieu A, Riglet L, Legrand J, Rozier F, Gaude T, Fobis-Loisy I. 2021. The molecular signatures of compatible and incompatible pollination in Arabidopsis. BMC Genomics 22:268.

58. Kolde R. 2019. pheatmap: Pretty Heatmaps. R package version 1.0. 12.

59. Kusaba M, Dwyer K, Hendershot J, Vrebalov J, Nasrallah JB, Nasrallah ME. 2001. Self-incompatibility in the genus Arabidopsis: characterization of the S locus in the outcrossing A. lyrata and its autogamous relative A. thaliana. Plant Cell 13:627–643.

60. Kusaba M, Tung C-W, Nasrallah ME, Nasrallah JB. 2002. Monoallelic expression and dominance interactions in anthers of self-incompatible Arabidopsis lyrata. Plant Physiol. 128:17–20.

61. Le Veve A, Burghgraeve N, Genete M, Lepers-Blassiau C, Takou M, De Meaux J, Mable BK, Durand E, Vekemans X, Castric V. 2022. Long-term balancing selection and the genetic load linked to the self-incompatibility locus in Arabidopsis halleri and A. lyrata. bioRxiv [Internet]:2022.04.12.487987. Available from: https://www.biorxiv.org/content/10.1101/2022.04.12.487987v1

62. Levin DA. 1975. Minority Cytotype Exclusion in Local Plant Populations. Taxon 24:35–43.

63. Levin DA. 2012. Mating system shifts on the trailing edge. Ann. Bot. 109:613–620.

64. Li H. 2018. Minimap2: pairwise alignment for nucleotide sequences. Bioinformatics 34:3094–3100.

65. Li H, Durbin R. 2009. Fast and accurate short read alignment with Burrows-Wheeler transform. Bioinformatics 25:1754–1760.

66. Li H, Handsaker B, Wysoker A, Fennell T, Ruan J, Homer N, Marth G, Abecasis G, Durbin R, 1000 Genome Project Data Processing Subgroup. 2009. The Sequence Alignment/Map format and SAMtools. Bioinformatics 25:2078–2079.

67. Li Y, van Kleunen M, Stift M. 2019. Genetic interaction between two unlinked loci underlies the loss of self-incompatibility in Arabidopsis lyrata. bioRxiv [Internet]:830414. Available from: https://www.biorxiv.org/content/10.1101/830414v1.full

68. Llaurens V, Billiard S, Leducq J-B, Castric V, Klein EK, Vekemans X. 2008. Does frequency-dependent selection with complex dominance interactions accurately predict allelic frequencies at the self-incompatibility locus in Arabidopsis halleri? Evolution 62:2545–2557.

69. Long Q, Rabanal FA, Meng D, Huber CD, Farlow A, Platzer A, Zhang Q, Vilhjálmsson BJ, Korte A, Nizhynska V, et al. 2013. Massive genomic variation and strong selection in Arabidopsis thaliana lines from Sweden. Nat. Genet. 45:884–890.

70. Lu Y. 2011. Arabidopsis Pollen Tube Aniline Blue Staining. Bio Protoc. [Internet] 1. Available from: https://bio-protocol.org/e88

71. Lysak MA, Berr A, Pecinka A, Schmidt R, McBreen K, Schubert I. 2006. Mechanisms of chromosome number reduction in Arabidopsis thaliana and related Brassicaceae species. Proc. Natl. Acad. Sci. U. S. A. 103:5224–5229.

72. Mable BK, Beland J, Di Berardo C. 2004. Inheritance and dominance of self-incompatibility alleles in polyploid Arabidopsis lyrata. Heredity 93:476–486.

73. Mable BK, Brysting AK, Jørgensen MH, Carbonell AKZ, Kiefer C, Ruiz-Duarte P, Lagesen K, Koch MA. 2018. Adding Complexity to Complexity: Gene Family Evolution in Polyploids. Frontiers in Ecology and Evolution [Internet] 6. Available from: https://www.frontiersin.org/articles/10.3389/fevo.2018.00114

74. Mable BK, Hagmann J, Kim S-T, Adam A, Kilbride E, Weigel D, Stift M. 2017. What causes mating system shifts in plants? Arabidopsis lyrata as a case study. Heredity 118:52–63.

75. Mable BK, Robertson AV, Dart S, Di Berardo C, Witham L. 2005. Breakdown of self-incompatibility in the perennial Arabidopsis lyrata (Brassicaceae) and its genetic consequences. Evolution 59:1437– 1448.

76. Mable BK, Schierup MH, Charlesworth D. 2003. Estimating the number, frequency, and dominance of S-alleles in a natural population of Arabidopsis lyrata(Brassicaceae) with sporophytic control of self-incompatibility. Heredity 90:422–431.

77. Mattila TM, Laenen B, Slotte T. 2020. Population genomics of transitions to selfing in Brassicaceae model systems. Statistical population genomics [Internet]. Available from: https://library.oapen.org/bitstream/handle/20.500.12657/23339/1006816.pdf?sequence=1#page=273

78. McGaugh SE, Noor MAF. 2012. Genomic impacts of chromosomal inversions in parapatric Drosophila species. Philos. Trans. R. Soc. Lond. B Biol. Sci. 367:422–429.

79. McKenna A, Hanna M, Banks E, Sivachenko A, Cibulskis K, Kernytsky A, Garimella K, Altshuler D, Gabriel S, Daly M, et al. 2010. The Genome Analysis Toolkit: a MapReduce framework for analyzing next-generation DNA sequencing data. Genome Res. 20:1297–1303.

80. Mishima M, Takayama S, Sasaki K, Jee JG, Kojima C, Isogai A, Shirakawa M. 2003. Structure of the male determinant factor for Brassica self-incompatibility. J. Biol. Chem. 278:36389–36395.

81. Nasrallah JB. 2019. Self-incompatibility in the Brassicaceae: Regulation and mechanism of self-recognition. Curr. Top. Dev. Biol. 131:435–452.

82. Nasrallah ME, Liu P, Sherman-Broyles S, Boggs NA, Nasrallah JB. 2004. Natural variation in expression of self-incompatibility in *Arabidopsis thaliana*: Implications for the evolution of selfing. Proceedings of the National Academy of Sciences 101:16070–16074.

83. Nei M. 1972. Genetic Distance between Populations. Am. Nat. 106:283–292.

84. Nordborg M, Donnelly P. 1997. The coalescent process with selfing. Genetics 146:1185–1195.

85. Novikova PY, Hohmann N, Nizhynska V, Tsuchimatsu T, Ali J, Muir G, Guggisberg A, Paape T, Schmid K, Fedorenko OM, et al. 2016. Sequencing of the genus Arabidopsis identifies a complex history of nonbifurcating speciation and abundant trans-specific polymorphism. Nat. Genet. 48:1077–1082.

86. Novikova PY, Kolesnikova UK, Scott AD. 2022. Ancestral self-compatibility facilitates the establishment of allopolyploids in Brassicaceae. Plant Reprod. [Internet]. Available from: http://dx.doi.org/10.1007/s00497-022-00451-6

87. Novikova PY, Tsuchimatsu T, Simon S, Nizhynska V, Voronin V, Burns R, Fedorenko OM, Holm S, Sall T, Prat E, et al. 2017. Genome Sequencing Reveals the Origin of the Allotetraploid Arabidopsis suecica. Mol. Biol. Evol. 34:957–968.

88. Okamoto S, Odashima M, Fujimoto R, Sato Y, Kitashiba H, Nishio T. 2007. Self-compatibility in Brassica napus is caused by independent mutations in S-locus genes. Plant J. 50:391–400.

89. Ortiz EM. 2019. vcf2phylip v2.0: convert a VCF matrix into several matrix formats for phylogenetic analysis. Available from: https://zenodo.org/record/2540861

90. Paape T, Briskine RV, Halstead-Nussloch G, Lischer HEL, Shimizu-Inatsugi R, Hatakeyama M, Tanaka K, Nishiyama T, Sabirov R, Sese J, et al. 2018. Patterns of polymorphism and selection in the subgenomes of the allopolyploid Arabidopsis kamchatica. Nat. Commun. 9:3909.

91. Paradis E, Claude J, Strimmer K. 2004. APE: Analyses of Phylogenetics and Evolution in R language. Bioinformatics 20:289–290.

92. Paradis E, Schliep K. 2019. ape 5.0: an environment for modern phylogenetics and evolutionary analyses in R. Bioinformatics 35:526–528.

93. Pembleton LW, Cogan NOI, Forster JW. 2013. StAMPP: an R package for calculation of genetic differentiation and structure of mixed-ploidy level populations. Mol. Ecol. Resour. 13:946–952.

94. Prigoda NL, Nassuth A, Mable BK. 2005. Phenotypic and genotypic expression of self-incompatibility haplotypes in Arabidopsis lyrata suggests unique origin of alleles in different dominance classes. Mol. Biol. Evol. 22:1609–1620.

95. Rawat V, Abdelsamad A, Pietzenuk B, Seymour DK, Koenig D, Weigel D, Pecinka A, Schneeberger K. 2015. Improving the Annotation of Arabidopsis lyrata Using RNA-Seq Data. PLoS One 10:e0137391.

96. Rieseberg LH. 2001. Chromosomal rearrangements and speciation. Trends Ecol. Evol. 16:351–358.

97. Robinson JT, Turner D, Durand NC, Thorvaldsdóttir H, Mesirov JP, Aiden EL. 2018. Juicebox.js Provides a Cloud-Based Visualization System for Hi-C Data. Cell Syst 6:256–258.e1.

98. Roux C, Pauwels M, Ruggiero MV, Charlesworth D, Castric V, Vekemans X. 2013. Recent and ancient signature of balancing selection around the S-locus in Arabidopsis halleri and A. lyrata. Mol. Biol. Evol. 30:435–447.

99. Rowan BA, Heavens D, Feuerborn TR, Tock AJ, Henderson IR, Weigel D. 2019. An Ultra High-Density Arabidopsis thaliana Crossover Map That Refines the Influences of Structural Variation and Epigenetic Features. Genetics 213:771–787.

100. Ruggiero MV, Jacquemin B, Castric V, Vekemans X. 2008. Hitch-hiking to a locus under balancing selection: high sequence diversity and low population subdivision at the S-locus genomic region in Arabidopsis halleri. Genet. Res. 90:37–46.

101. Sarret G, Willems G, Isaure M-P, Marcus MA, Fakra SC, Frérot H, Pairis S, Geoffroy N, Manceau A, Saumitou-Laprade P. 2009. Zinc distribution and speciation in Arabidopsis halleri x Arabidopsis lyrata progenies presenting various zinc accumulation capacities. New Phytol. 184:581–595.

102. Schierup MH, Mable BK, Awadalla P, Charlesworth D. 2001. Identification and characterization of a polymorphic receptor kinase gene linked to the self-incompatibility locus of Arabidopsis lyrata. Genetics 158:387–399.

103. Schierup MH, Vekemans X, Christiansen FB. 1997. Evolutionary dynamics of sporophytic self-incompatibility alleles in plants. Genetics 147:835–846.

104. Schmickl R, Jorgensen MH, Brysting AK, Koch MA. 2010. The evolutionary history of the Arabidopsis lyrata complex: a hybrid in the amphi-Beringian area closes a large distribution gap and builds up a genetic barrier. BMC Evol. Biol. 10:98.

105. Schopfer CR, Nasrallah ME, Nasrallah JB. 1999. The male determinant of self-incompatibility in Brassica. Science 286:1697–1700.

106. Seppey M, Manni M, Zdobnov EM. 2019. BUSCO: Assessing Genome Assembly and Annotation Completeness. In: Kollmar M, editor. Gene Prediction: Methods and Protocols. New York, NY: Springer New York. p. 227–245.

107. Seregin A. 2023. Moscow University Herbarium (MW). Available from: http://dx.doi.org/10.15468/CPNHCC

108. Sherman-Broyles S, Boggs N, Farkas A, Liu P, Vrebalov J, Nasrallah ME, Nasrallah JB. 2007. S Locus Genes and the Evolution of Self-Fertility in Arabidopsis thaliana. Plant Cell 19:94.

109. Shiba H, Kakizaki T, Iwano M, Tarutani Y, Watanabe M, Isogai A, Takayama S. 2006. Dominance relationships between self-incompatibility alleles controlled by DNA methylation. Nat. Genet. 38:297–299.

110. Shimizu-Inatsugi R, Lihová J, Iwanaga H, Kudoh H, Marhold K, Savolainen O, Watanabe K, Yakubov VV, Shimizu KK. 2009. The allopolyploid Arabidopsis kamchatica originated from multiple individuals of Arabidopsis lyrata and Arabidopsis halleri. Mol. Ecol. 18:4024–4048.

111. Shimizu KK, Fujii S, Marhold K, Watanabe K, Kudoh H. 2005. Arabidopsis kamchatica (Fisch. ex DC.) К. Shimizu & Kudoh and A. kamchatica subsp. kawasakiana (Makino) K. Shimizu & Kudoh, New Combinations. Acta phytotaxonomica et geobotanica 56:163–172.

112. Shimizu KK, Tsuchimatsu T. 2015. Evolution of Selfing: Recurrent Patterns in Molecular Adaptation. Annu. Rev. Ecol. Evol. Syst. 46:593–622.

113. Shumate A, Salzberg SL. 2020. Liftoff: accurate mapping of gene annotations. Bioinformatics [Internet]. Available from: http://dx.doi.org/10.1093/bioinformatics/btaa1016

114. Sicard A, Lenhard M. 2011. The selfing syndrome: a model for studying the genetic and evolutionary basis of morphological adaptation in plants. Ann. Bot. 107:1433–1443.

115. Slotte T, Hazzouri KM, Ågren JA, Koenig D, Maumus F, Guo Y-L, Steige K, Platts AE, Escobar JS, Newman LK, et al. 2013. The Capsella rubella genome and the genomic consequences of rapid mating system evolution. Nat. Genet. 45:831–835.

116. Smit AFA, Hubley R. 2008-2015. RepeatModeler Open-1.0 http://www.repeatmasker.org.

117. Smit AFA, Hubley R, Green P. 2013-2015. RepeatMasker Open-4.0. .

118. Stamatakis A. 2014. RAxML version 8: a tool for phylogenetic analysis and post-analysis of large phylogenies. Bioinformatics 30:1312–1313.

119. Stein JC, Howlett B, Boyes DC, Nasrallah ME, Nasrallah JB. 1991. Molecular cloning of a putative receptor protein kinase gene encoded at the self-incompatibility locus of Brassica oleracea. Proc. Natl. Acad. Sci. U. S. A. 88:8816–8820.

120. Stevison LS, Hoehn KB, Noor MAF. 2011. Effects of Inversions on Within- and Between-Species Recombination and Divergence. Genome Biol. Evol. 3:830–841.

121. Takayama S, Isogai A. 2005. Self-incompatibility in plants. Annu. Rev. Plant Biol. 56:467–489.

122. Takayama S, Shiba H, Iwano M, Shimosato H, Che FS, Kai N, Watanabe M, Suzuki G, Hinata K, Isogai A. 2000. The pollen determinant of self-incompatibility in *Brassica campestris*. Proc. Natl. Acad. Sci. U. S. A. 97:1920–1925.

123. Takayama S, Shimosato H, Shiba H, Funato M, Che FS, Watanabe M, Iwano M, Isogai A. 2001. Direct ligand-receptor complex interaction controls Brassica self-incompatibility. Nature 413:534–538.

124. Takou M, Hämälä T, Koch EM, Steige KA, Dittberner H, Yant L, Genete M, Sunyaev S, Castric V, Vekemans X, et al. 2021. Maintenance of Adaptive Dynamics and No Detectable Load in a Range-Edge Outcrossing Plant Population. Mol. Biol. Evol. 38:1820–1836.

125. Takou M, Hämälä T, Koch E, Steige KA, Dittberner H, Yant L, Genete M, Sunyaev S, Castric V, Vekemans X, et al. 2020. Maintenance of adaptive dynamics and no detectable load in a range-edge out-crossing plant population. bioRxiv [Internet]:709873. Available from: https://www.biorxiv.org/content/10.1101/709873v3

126. Tarutani Y, Shiba H, Iwano M, Kakizaki T, Suzuki G, Watanabe M, Isogai A, Takayama S. 2010. Trans-acting small RNA determines dominance relationships in Brassica self-incompatibility. Nature 466:983–986.

127. The 1001 Genomes Consortium. 2016. 1,135 Genomes Reveal the Global Pattern of Polymorphism in Arabidopsis thaliana. Cell 166:481–491.

128. Tsuchimatsu T, Fujii S. 2022. The selfing syndrome and beyond: diverse evolutionary consequences of mating system transitions in plants. Philos. Trans. R. Soc. Lond. B Biol. Sci. 377:20200510.

129. Tsuchimatsu T, Goubet PM, Gallina S, Holl A-C, Fobis-Loisy I, Bergès H, Marande W, Prat E, Meng D, Long Q, et al. 2017. Patterns of Polymorphism at the Self-Incompatibility Locus in 1,083 Arabidopsis thaliana Genomes. Mol. Biol. Evol. 34:1878–1889.

130. Tsuchimatsu T, Kaiser P, Yew CL, Bachelier JB, Shimizu KK. 2012. Recent loss of self-incompatibility by degradation of the male component in allotetraploid Arabidopsis kamchatica. PLoS Genet. 8:e1002838.

131. Tsuchimatsu T, Shimizu KK. 2013. Effects of pollen availability and the mutation bias on the fixation of mutations disabling the male specificity of self-incompatibility. J. Evol. Biol. 26:2221–2232.

132. Tsuchimatsu T, Suwabe K, Shimizu-Inatsugi R, Isokawa S, Pavlidis P, Städler T, Suzuki G, Takayama S, Watanabe M, Shimizu KK. 2010. Evolution of self-compatibility in Arabidopsis by a mutation in the male specificity gene. Nature 464:1342–1346.

133. Uyenoyama MK, Zhang Y, Newbigin E. 2001. On the origin of self-incompatibility haplotypes: transition through self-compatible intermediates. Genetics 157:1805–1817.

134. Vekemans X, Poux C, Goubet PM, Castric V. 2014. The evolution of selfing from outcrossing ancestors in Brassicaceae: what have we learned from variation at the S-locus? J. Evol. Biol. 27:1372–1385.

135. Vekemans X, Slatkin M. 1994. Gene and allelic genealogies at a gametophytic self-incompatibility locus. Genetics 137:1157–1165.

136. Waterhouse AM, Procter JB, Martin DM, Clamp M, Barton GJ. 2009. Jalview Version 2--a multiple sequence alignment editor and analysis workbench. Bioinformatics 25:1189–1191.

137. Willi Y, Lucek K, Bachmann O, Walden N. 2022. Recent speciation associated with range expansion and a shift to self-fertilization in North American Arabidopsis. Nat. Commun. 13:7564.

138. Willi Y, Määttänen K. 2010. Evolutionary dynamics of mating system shifts in Arabidopsis lyrata. J. Evol. Biol. 23:2123–2131.

139. Wingett S, Ewels P, Furlan-Magaril M, Nagano T, Schoenfelder S, Fraser P, Andrews S. 2015. HiCUP: pipeline for mapping and processing Hi-C data. F1000Res. 4:1310.

140. Wolff J, Rabbani L, Gilsbach R, Richard G, Manke T, Backofen R, Grüning BA. 2020. Galaxy HiCExplorer 3: a web server for reproducible Hi-C, capture Hi-C and single-cell Hi-C data analysis, quality control and visualization. Nucleic Acids Res. 48:W177–W184.

141. Wright S. 1939. The Distribution of Self-Sterility Alleles in Populations. Genetics 24:538–552.

142. Yu G. 2020. Using ggtree to Visualize Data on Tree-Like Structures. Curr. Protoc. Bioinformatics 69:e96.

143. Yu G, Lam TT-Y, Zhu H, Guan Y. 2018. Two Methods for Mapping and Visualizing Associated Data on Phylogeny Using Ggtree. Mol. Biol. Evol. 35:3041–3043.

144. Yu G, Smith DK, Zhu H, Guan Y, Lam TT-Y. 2017. Ggtree : An r package for visualization and annotation of phylogenetic trees with their covariates and other associated data. Methods Ecol. Evol. 8:28–36.

145. Zhang T, Qiao Q, Novikova PY, Wang Q, Yue J, Guan Y, Ming S, Liu T, De J, Liu Y, et al. 2019. Genome of Crucihimalaya himalaica, a close relative of Arabidopsis, shows ecological adaptation to high altitude. Proc. Natl. Acad. Sci. U. S. A. 116:7137–7146.

146. Zhao H, Zhang Y, Zhang H, Song Y, Zhao F, Zhang Y ’e, Zhu S, Zhang H, Zhou Z, Guo H, et al. 2022. Origin, loss, and regain of self-incompatibility in angiosperms. Plant Cell 34:579–596.

147. Zhu W, Hu B, Becker C, Doğan ES, Berendzen KW, Weigel D, Liu C. 2017. Altered chromatin compaction and histone methylation drive non-additive gene expression in an interspecific Arabidopsis hybrid. Genome Biol. 18:157.

