## Supplementary Materials for "Transition to self-compatibility associated with dominant *S*-allele in a diploid Siberian progenitor of allotetraploid *Arabidopsis kamchatica* revealed by *Arabidopsis lyrata* genomes"

Supplementary Table 1. List of sequenced and analyzed *A. lyrata* accessions

Supplementary Table 2. List of all possible inversions between *A. lyrata* NT1, MN47v1, MN47v2, *A. suecica* and *C. rubella*.

Supplementary Table 3. Inversions between *A. lyrata* NT1 and MN47v2, as reported by CoGe Synmap after confirmation using Hi-C and examination of long reads mapping (see Supplementary Fig. 3-7).

| ***A. lyrata* - MN47 v2** | **start** | **end** | **length** | ***A. lyrata* - NT1** | **start** | **end** | **length** |
|---|---|---|---|---|---|---|---|
| chr 1 | 14625309 | 17069605 | 2444296 | chr 1 | 17471724 | 14410341 | 3061383 |
| chr 3 | 22131279 | 24228593 | 2097314 | chr 3 | 26447300 | 24296296 | 2151004 |
| chr 4 | 18459828 | 18815810 | 355982 | chr 4 | 19135397 | 18871811 | 263586 |
| chr 6 | 17833645 | 19172561 | 1338916 | chr 6 | 19669769 | 18509483 | 1160286 |
| chr 2 | 4890607 | 5375291 | 484684 | chr 2 | 4823531 | 4397789 | 425742 |

Supplementary Table 4. Demographic parameter estimates for all tested models.
Models correspond to those shown in Supplementary Figure 14. Parameters estimated by fastsimcoal26. Model with lowest AIC score highlighted in gray, with 95% confidence interval shown. Additional details (.tpl and .est files) at https://github.com/novikovalab/selfing_Alyrata.

| | Parameter estimates | | | | | | | | | | | | | |
|---|---|---|---|---|---|---|---|---|---|---|---|---|---|---|
| Model | **NPOP1** | **NPOP2** | **NANC** | **TDIV** | **migr12** | **migr21** | | | | | **MaxEstLhood** | **deltaL** | **AIC** | **AIC weight** |
| A | 9,019 | 94,077 | 62,285 | 29,166 | - | - | | | | | -4443296.453 | 314148.808 | 20462143.5 | 0 |
| B | 7,155 | 100,657 | 62 | 103,935 | 1.76E-06 | - | | | | | -4434947.57 | 305799.925 | 20423697.47 | 0 |
| C | 5,216 | 95,938 | 25,077 | 79,794 | 1.11E-06 | 5.19E-06 | | | | | -4431951.327 | 302803.682 | 20409901.27 | 0 |
| | **TWO_NE_PRESENT_POP1** | **TWO_NE_PRESENT_POP2** | **TWO_NE_ANC** | **T_SPL** | **migr12** | **migr21** | **RESIZE_ANC** | **SEVERITY_BOT** | **T_BOT_POP1_BEG** | **TWO_NE_BOT_POP1** | **MaxEstLhood** | **deltaL** | **AIC** | **AIC weight** |
| D | 508,491 | 95,228 | 330,000 | 31,864 | - | - | 0.649 | 2.641 | 2952 | 378 | -4436717.142 | 307569.497 | 20431848.65 | 0 |
| E | 1,052,979 | 66,800 | 258,485 | 87,756 | 8.05E-06 | 7.21E-06 | 0.245 | 3.681 | 2034 | 27 | -4420925.706 | 291778.061 | 20359130.41 | 1 |
| 2.5% | 11,203 | 40,447 | 2,836 | 36,002 | 3.76E-09 | 1.49E-09 | 0.135 | 1.283 | 1337.45 | 10 | - | - | - | - |
| 97.5% | 1,105,516 | 113,109 | 551,273 | 89,150 | 2.42E-05 | 2.01E-05 | 0.694 | 9.468 | 6809.1 | 96.625 | - | - | - | - |

Supplementary Data 1: Multisequence fasta of SRK sequences used in genotyping

Supplementary Data 2: Multisequence fasta of SCR sequences used in genotyping

Supplementary Data 3: Alignment for first exon of SRK used for phylogenetic inference

Supplementary Data 4: Maximum likelihood phylogeny in Newick format (lyrata.tre)

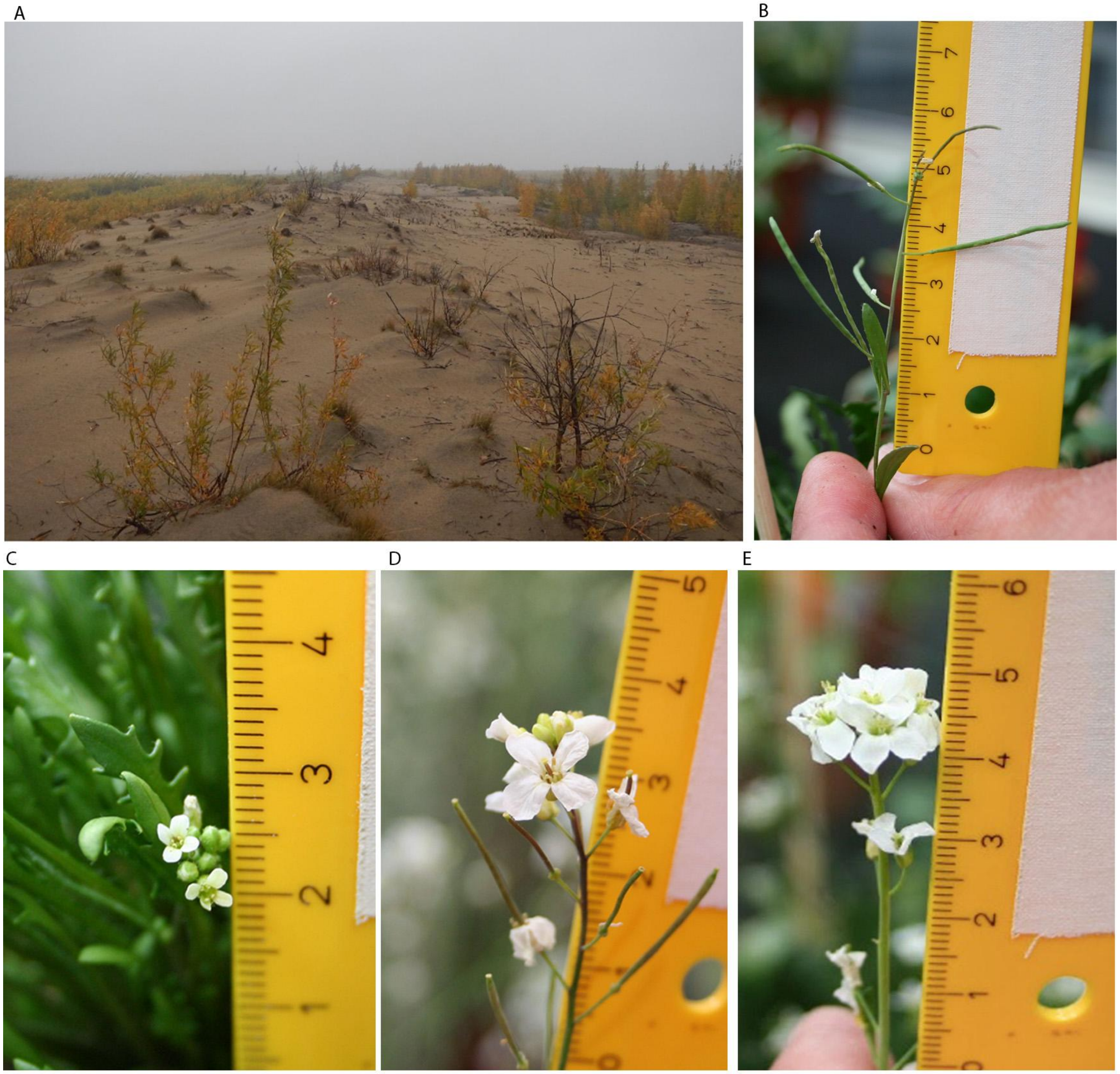

Supplementary Figure 1. (A) Photo of the collection site of the NT1 accession, showing the natural habitat of *A. lyrata* on an island in the course of the Lena River. (B) Long fruits formed by NT1 *A. lyrata* accession in the green house without artificial cross-fertilisation. (C) Flowers of NT1 *A. lyrata* Siberian accession appear to be smaller in size compared to (D) North American selfing MN47 and (E) outcrossing *A. lyrata* accessions growing in the greenhouse.

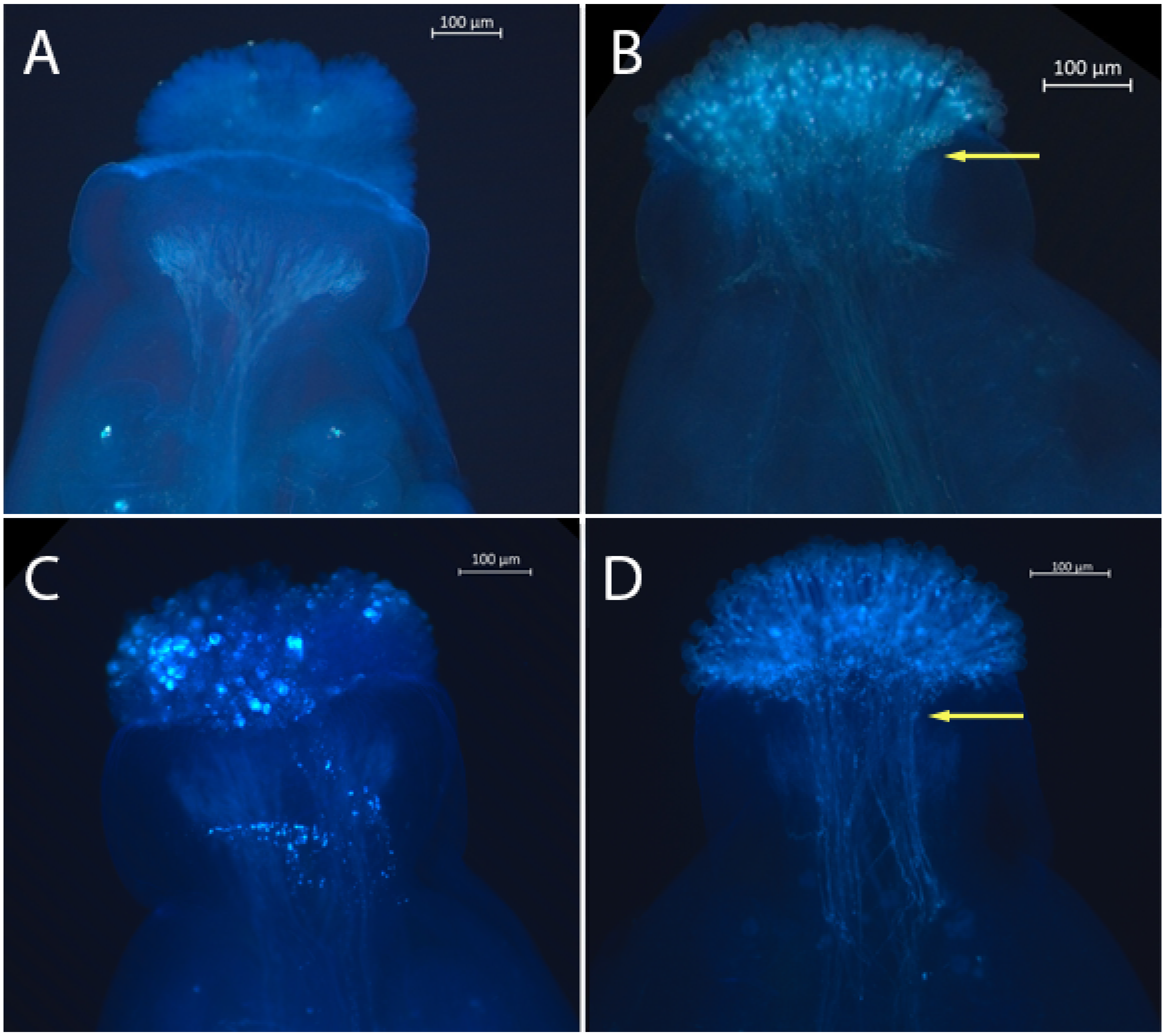

Supplementary Figure 2. Pollen tube staining confirming self-compatibility of the NT1. (A) Unpollinated stigma. (B) Self-compatible reaction: pollen tube growth on stigma of *A. lyrata* NT1 flower after self-pollination. (C) Self-incompatible reaction in outcrossing accession after self-pollination: pollen on the top of the stigma. (D) Compatible reaction: pollen tube growth in outcrossing *A. lyrata* accession after cross-pollination.

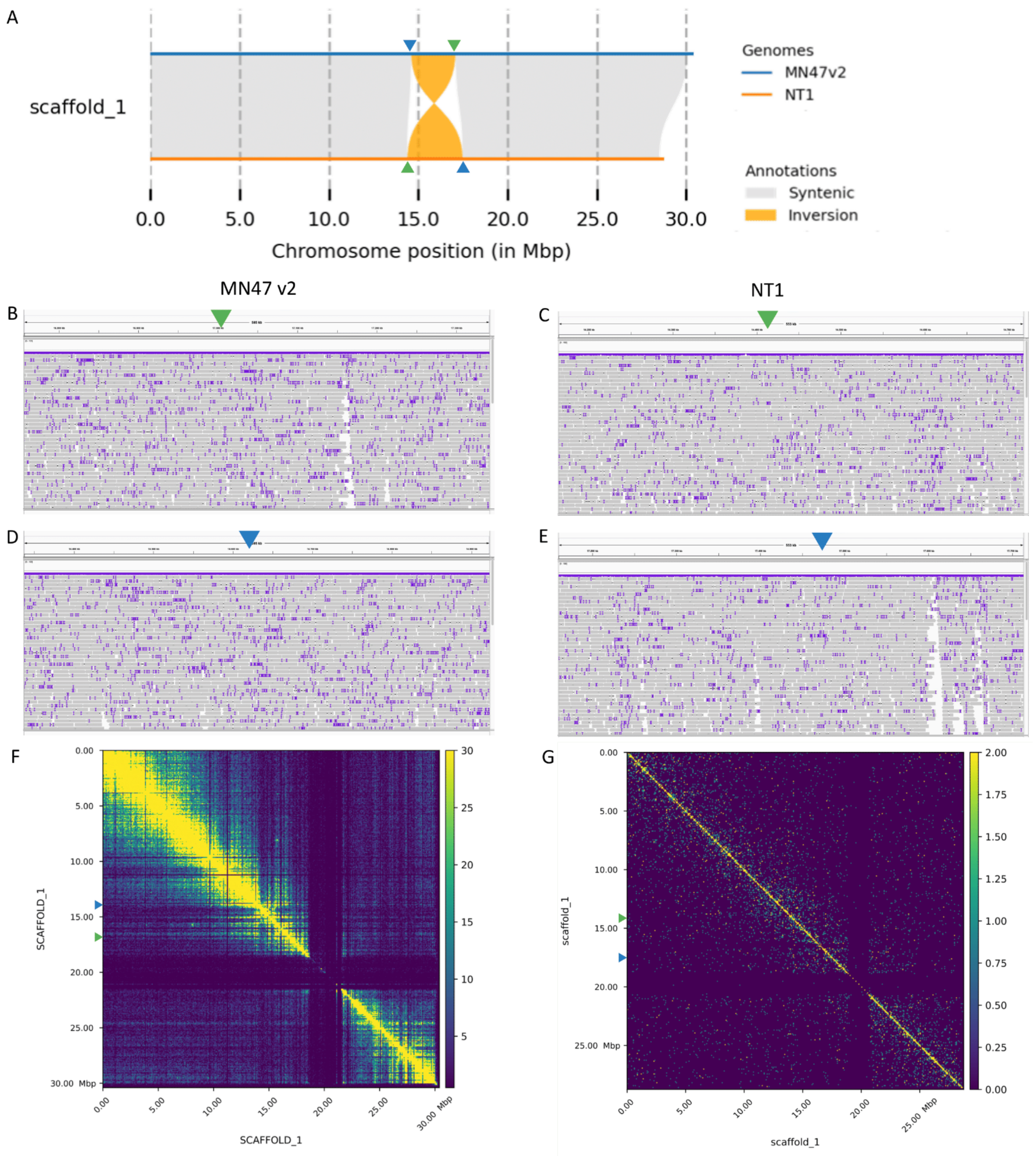

Supplementary Figure 3. Validation of an inversion on chromosome 1 between *A. lyrata* NT1 and MN47 accessions. a) Synteny and rearrangement plot of chromosome 1. (b,d) IGV snapshot showing PacBio HiFi reads of MN47 mapped to MN47 v2 assembly around inversion breakpoints (Supplementary Table 1, first row). (c,e) IGV snapshot showing PacBio HiFi reads of NT1 mapped to the NT1 assembly around inversion breakpoints (Supplementary Table 1, first row). The continuity of the long reads mapped in (b-e) suggests the inferred inversion is real. The results of the Hi-C contact maps of MN47 (f) and NT1 (g) show that a majority of the paired reads are mapped in cis, again validating the inversion. The colors on the Hi-C heatmaps (f,g) correspond to the density of the paired reads from low (dark blue) to high (yellow). The triangle markers in (a-g) indicate the start (blue triangle) and the stop (green triangle) of inversion.

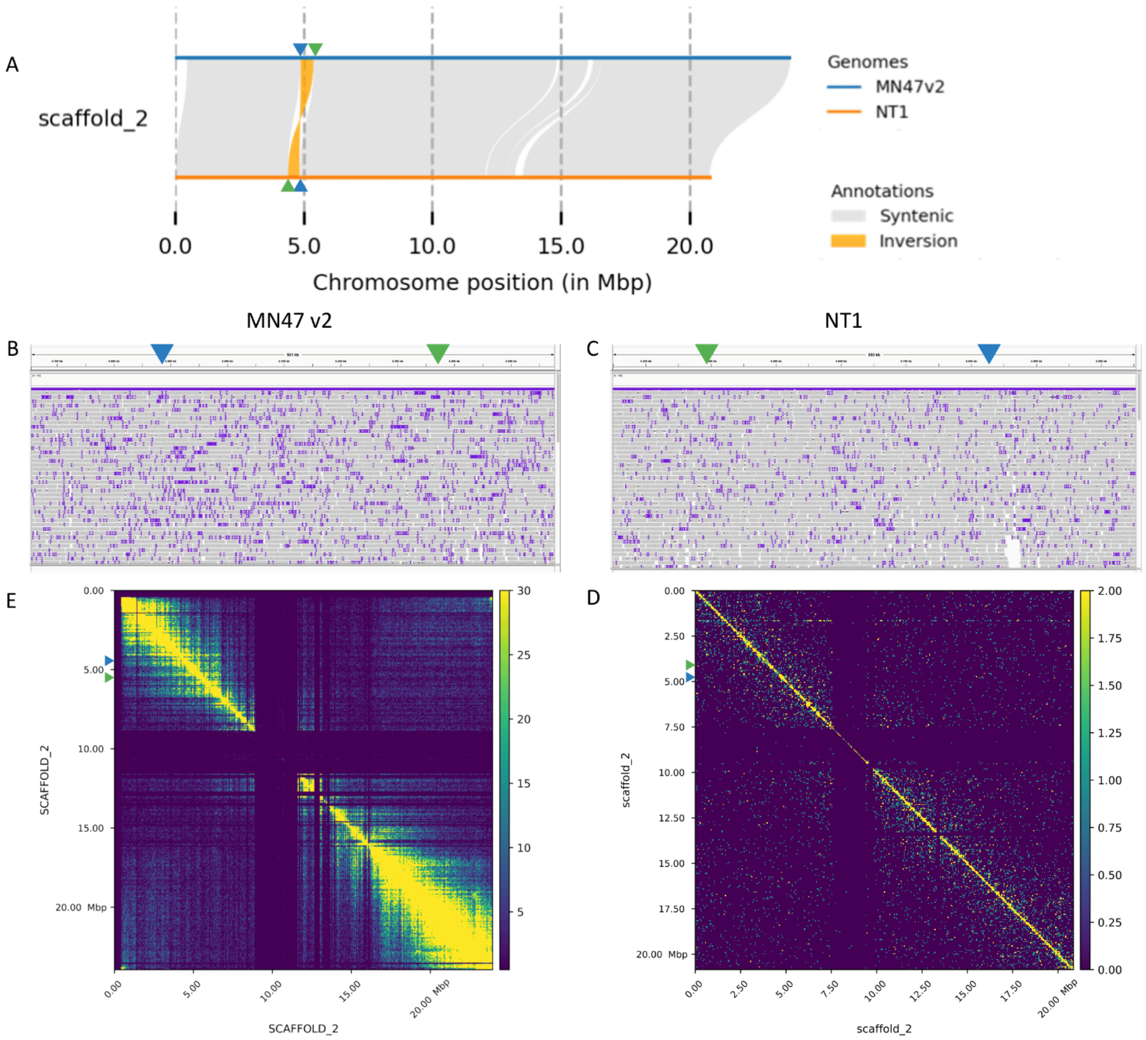

Supplementary Figure 4. Validation of an inversion on chromosome 2 between *A. lyrata* NT1 and MN47 accessions. a) Synteny and rearrangement plot of chromosome 2. b) IGV snapshot showing PacBio HiFi reads of MN47 mapped to MN47 v2 assembly around inversion breakpoints (Supplementary Table 1, second row). c) IGV snapshot showing PacBio HiFi reads of NT1 mapped to the NT1 assembly around inversion breakpoints (Supplementary Table 1, second row). The continuity of the long reads mapped in (b-c) suggests the inferred inversion is real. The results of the Hi-C contact maps of MN47 (d) and NT1 (e) show that a majority of the paired reads are mapped in cis, again validating the inversion. The colors on the Hi-C heatmaps (d,e) correspond to the density of the paired reads from low (dark blue) to high (yellow). The triangle markers in (a-e) indicate the start (blue triangle) and the stop (green triangle) of inversion.

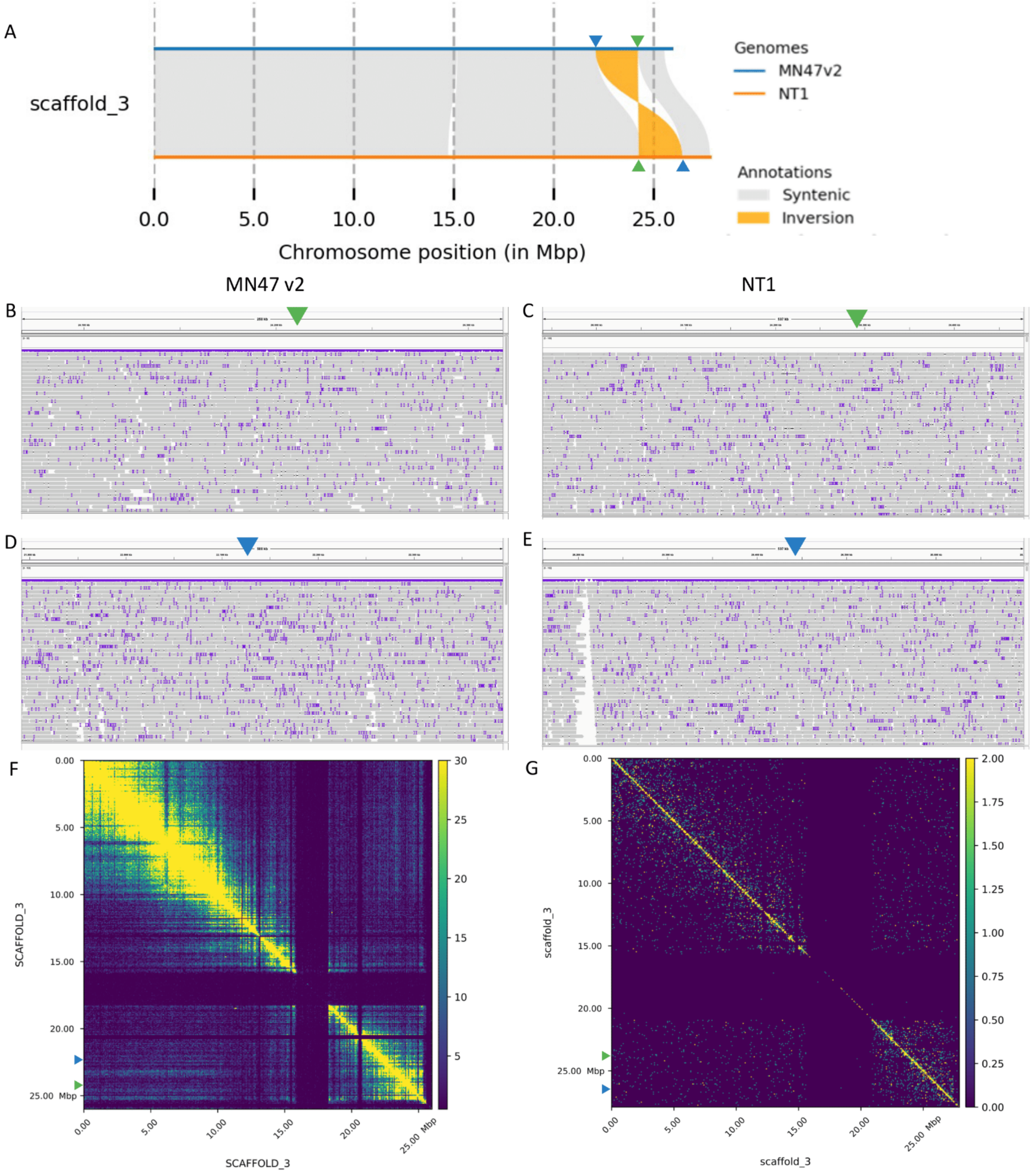

Supplementary Figure 5. Validation of an inversion on chromosome 3 between *A. lyrata* NT1 and MN47 accessions. a) Synteny and rearrangement plot of chromosome 3. (b,d) IGV snapshot showing PacBio HiFi reads of MN47 mapped to MN47 v2 assembly around inversion breakpoints (Supplementary Table 1, third row). (c,e) IGV snapshot showing PacBio HiFi reads of NT1 mapped to the NT1 assembly around inversion breakpoints (Supplementary Table 1, third row). The continuity of the long reads mapped in (b-e) suggests the inferred inversion is real. The results of the Hi-C contact maps of MN47 (f) and NT1 (g) show that a majority of the paired reads are mapped in cis, again validating the inversion. The colors on the Hi-C heatmaps (f,g) correspond to the density of the paired reads from low (dark blue) to high (yellow). The triangle markers in (a-g) indicate the start (blue triangle) and the stop (green triangle) of inversion.

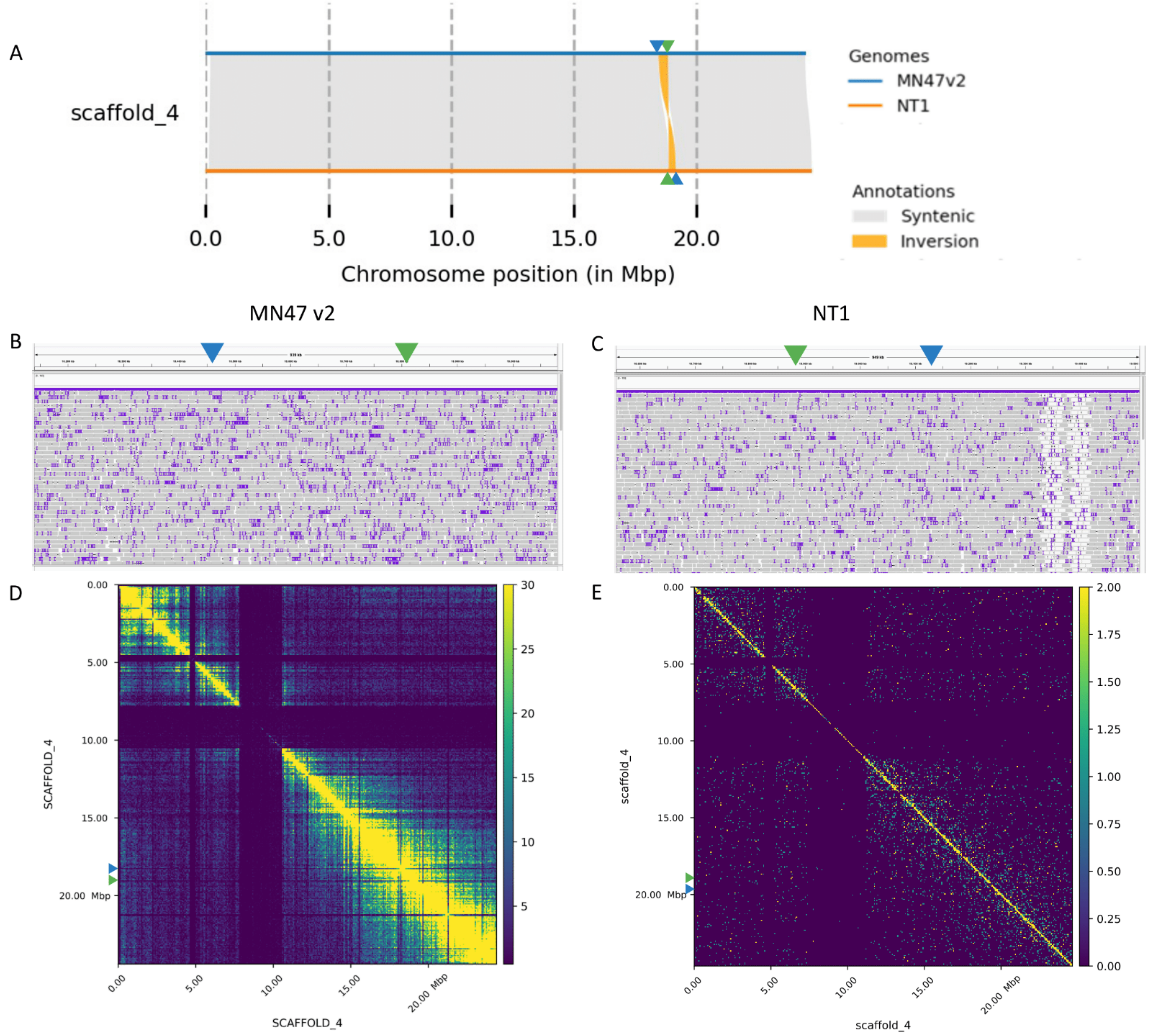

Supplementary Figure 6. Validation of an inversion on chromosome 4 between *A. lyrata* NT1 and MN47 accessions. a) Synteny and rearrangement plot of chromosome 4. b) IGV snapshot showing PacBio HiFi reads of MN47 mapped to MN47 v2 assembly around inversion breakpoints (Supplementary Table 1, fourth row). c) IGV snapshot showing PacBio HiFi reads of NT1 mapped to the NT1 assembly around inversion breakpoints (Supplementary Table 1, fourth row). The continuity of the long reads mapped in (b-c) suggests the inferred inversion is real. The results of the Hi-C contact maps of MN47 (d) and NT1 (e) show that a majority of the paired reads are mapped in cis, again validating the inversion. The colors on the Hi-C heatmaps (d,e) correspond to the density of the paired reads from low (dark blue) to high (yellow). The triangle markers in (a-e) indicate the start (blue triangle) and the stop (green triangle) of inversion.

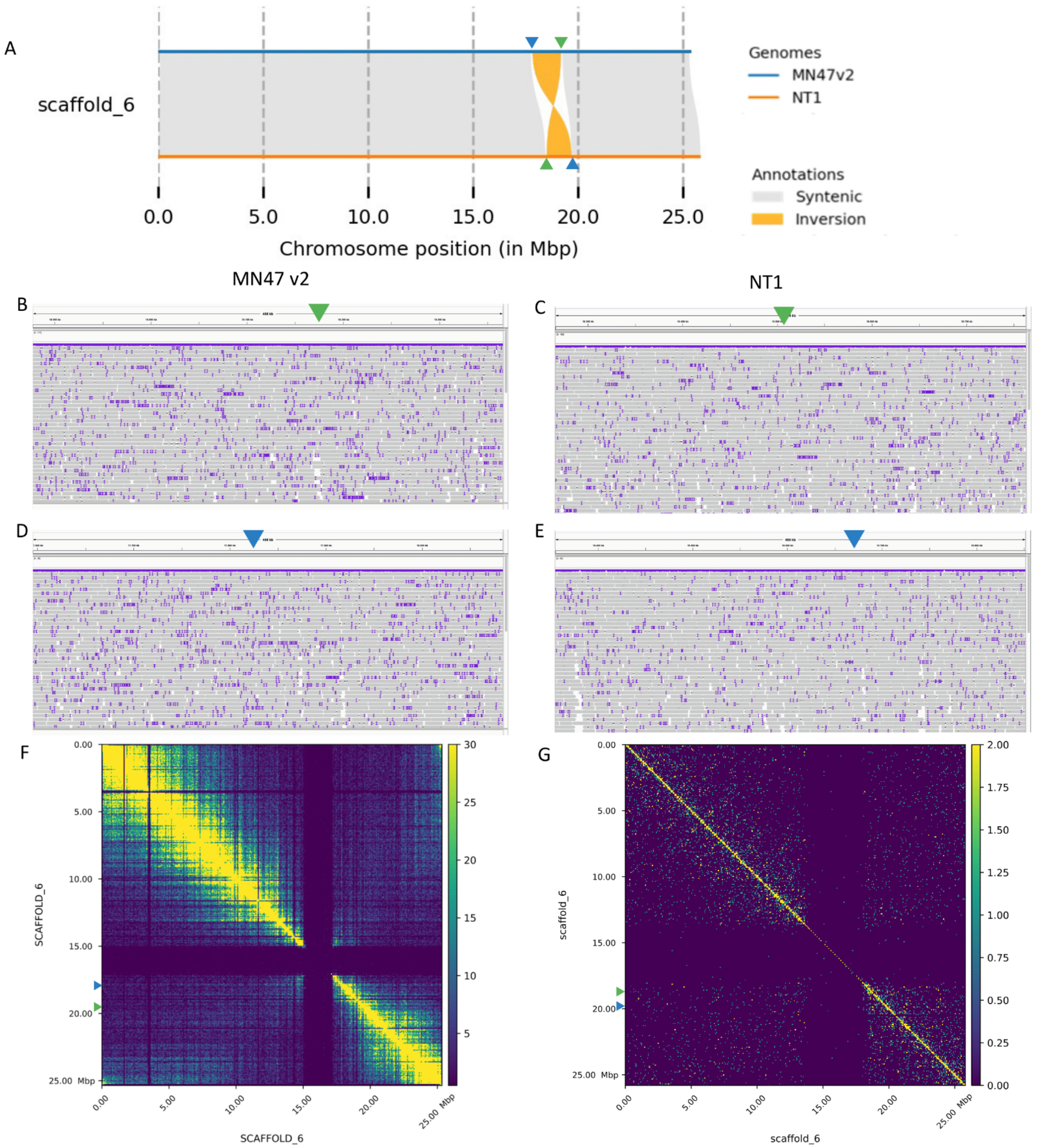

Supplementary Figure 7. Validation of an inversion on chromosome 6 between *A. lyrata* NT1 and MN47 accessions. a) Synteny and rearrangement plot of chromosome 6. (b,d) IGV snapshot showing PacBio HiFi reads of MN47 mapped to MN47 v2 assembly around inversion breakpoints (Supplementary Table 1, fifth row). (c,e) IGV snapshot showing PacBio HiFi reads of NT1 mapped to the NT1 assembly around inversion breakpoints (Supplementary Table 1, fifth row). The continuity of the long reads mapped in (b-e) suggests the inferred inversion is real. The results of the Hi-C contact maps of MN47 (f) and NT1 (g) show that a majority of the paired reads are mapped in cis, again validating the inversion. The colors on the Hi-C heatmaps (f,g) correspond to the density of the paired reads from low (dark blue) to high (yellow). The triangle markers in (a-g) indicate the start (blue triangle) and the stop (green triangle) of inversion.

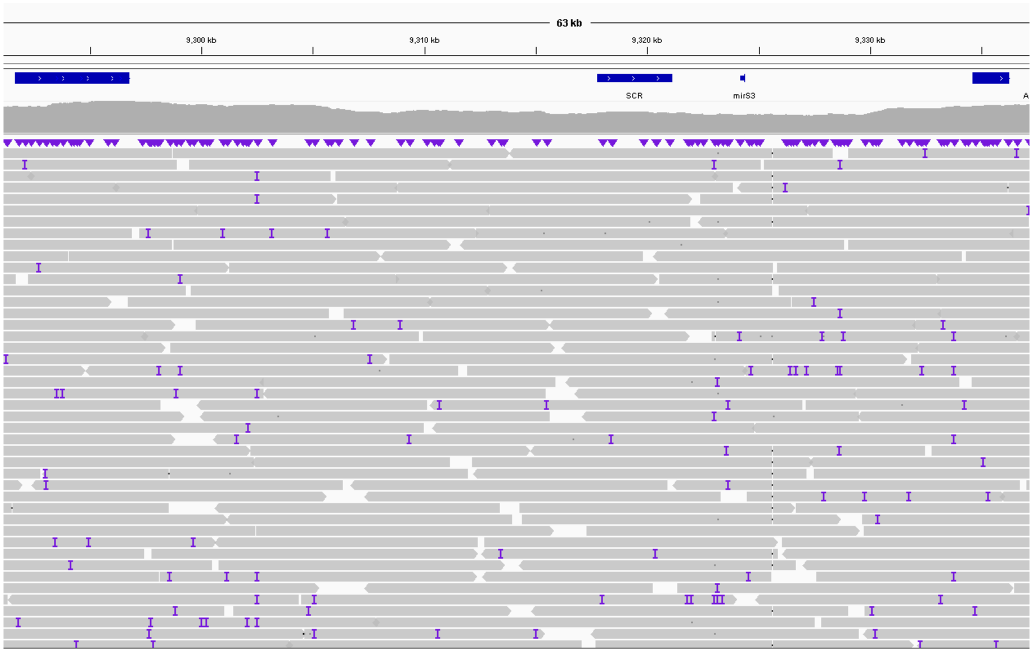

Supplementary Figure 8. IGV (Robinson et al. 2011) snapshot showing raw PacBio HiFi long-read coverage along the assembled *S*-locus in NT1 *A. lyrata* accession, on Scaffold 7. ARK3 and U-box flanking genes are marking the two ends of the *S*-locus, SCR gene is present, SRK is missing, mirS3 precursor is annotated manually. No gaps in the long read mapping suggests the completeness of the *S*-locus assembly.

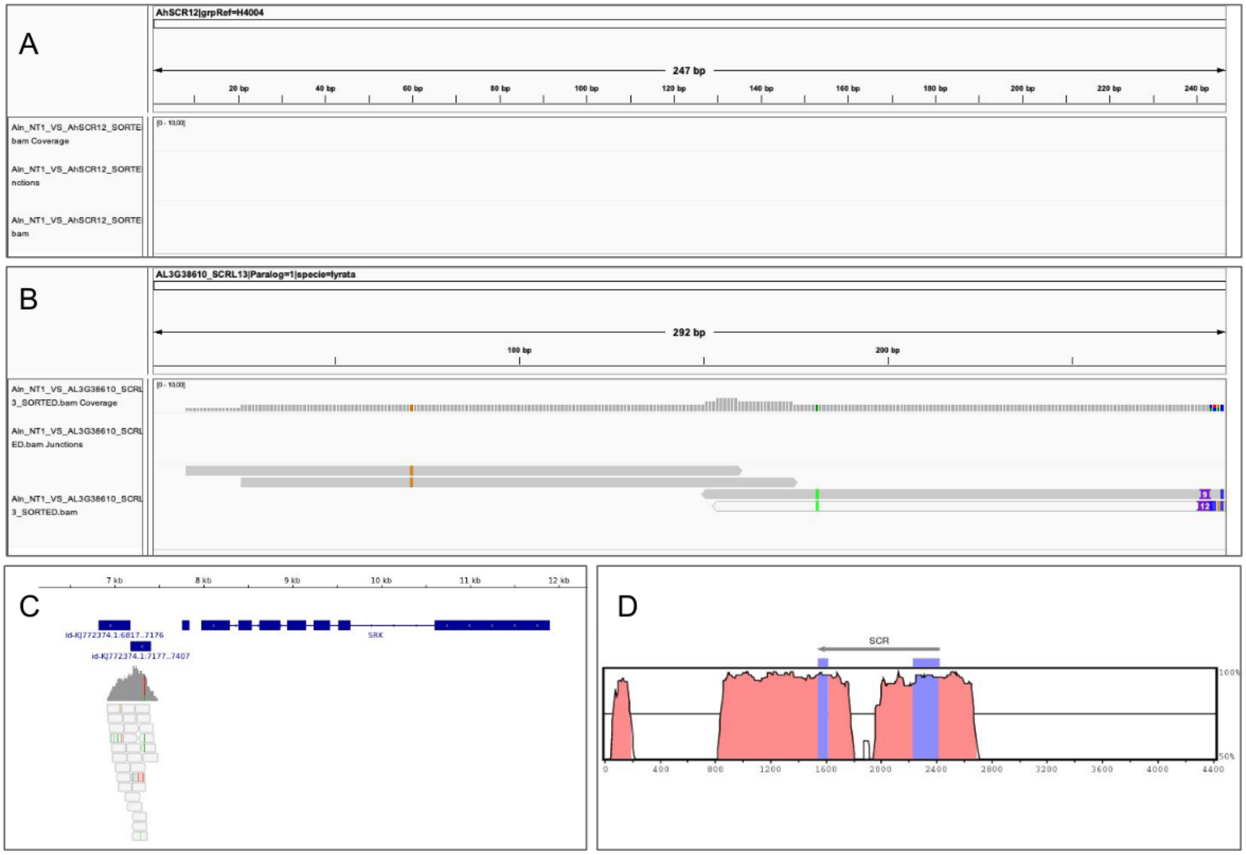

Supplementary Figure 9. IGV snapshot of the *A. lyrata* NT1 short reads from RNA-seq on flowers mapping to (A) the AhS12 SCR sequence, (B) the AL3G38610 SCR paralog, serving as positive control. The results suggest absence of expression of the SCR gene in NT1, although insufficient sequencing depth or transient expression in another stage of flower developmen cannot be ruled out. (C) IGV snapshot of the *A. lyrata* NT1 short reads mapping to the combined reference of NT1 genome (with absent SRK) and AhS12 sequence with AhSRK12. The absence of reads mapping to AhSRK12 confirms the loss of SRK from the NT1 *A. lyrata* genome. (D) Vista plot comparing the SCR region within the S-locus of Ah12 from *Arabidopsis halleri* (used as a reference) with that of NT1. Strong similarity is observed in the promoter region suggesting the absence of major rearrangements that could have led to loss of expression.

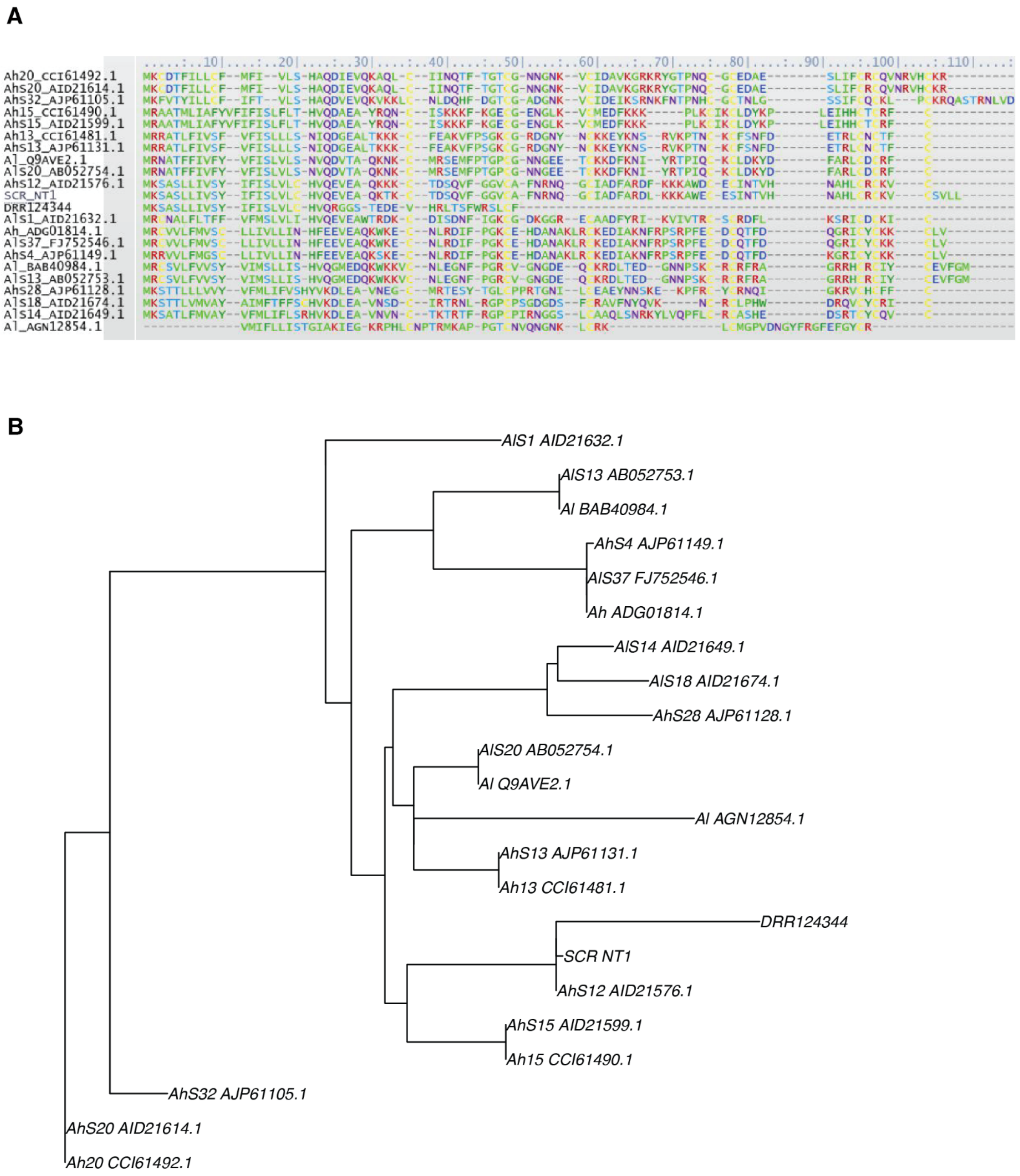

Supplementary Fig. 10. (A) Protein alignment of SCR including previously published sequences and SCR sequences from NT1 and DRR124344 (lyrpet4). SCR from lyrpet4 appears to be shorter and degraded compared to the rest, which may explain why the genotyping pipeline fails to identify it. (B) The maximum likelihood tree shows that DRR124344 (lyrpet4) clusters together with NT1 and AhS12.

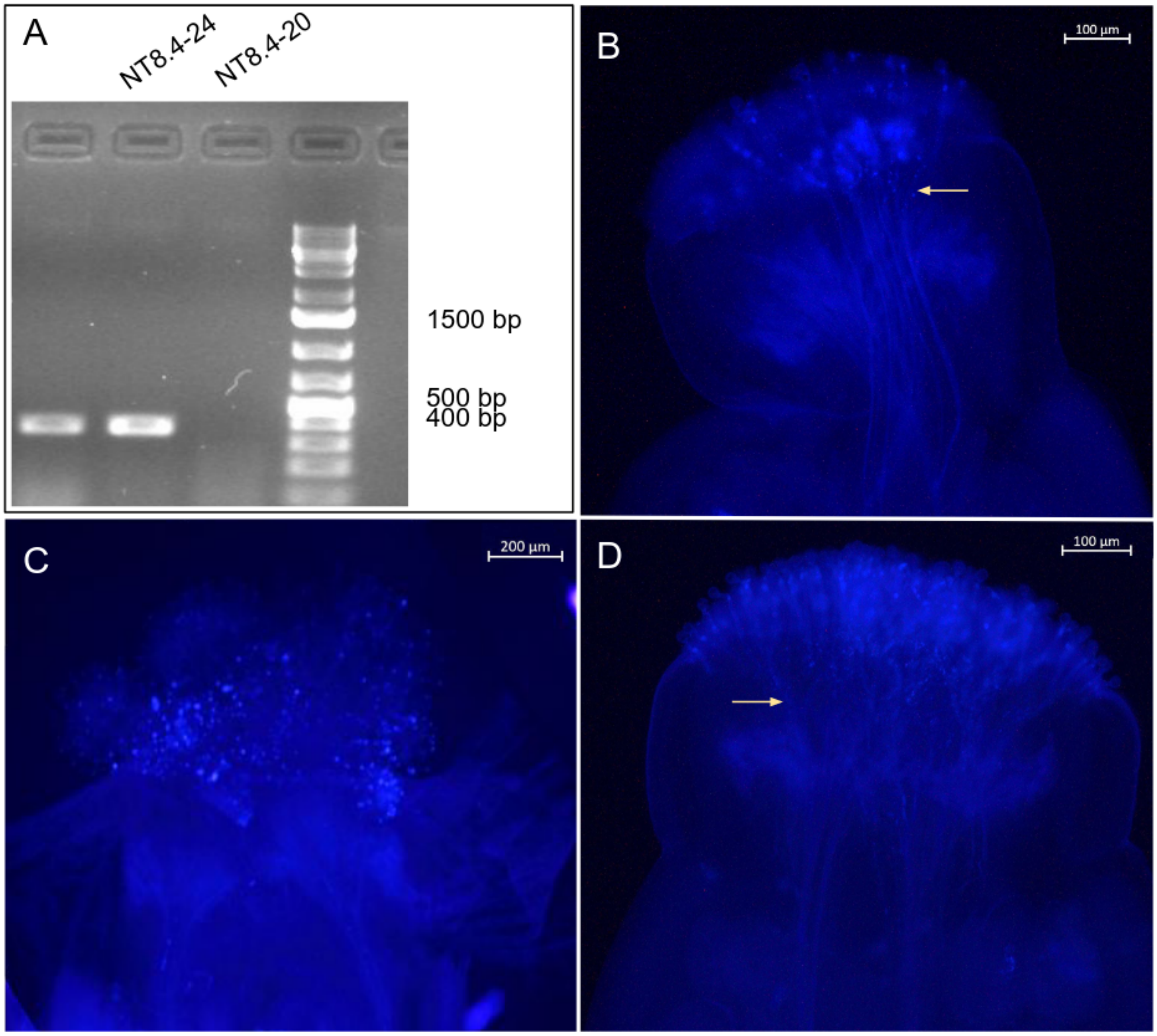

Supplementary figure 11. (A) Gel picture with PCR products and 1kb+ ladder identifying accessions with AhSRK12 (two positive bands with expected size of 360 bp). The sample in the middle (NT8.4-24) was used for crosses with NT1 (B) and another outcrossing accession without AhSRK12 (on the right, NT8.4-20) was used in (D). (B) NT8.4-24 (SRK12) ♀ x NT1 (SCR12) ♂ = compatible, yellow arrows show the growing pollen tubes. This suggests that SCR12 in NT1 is not recognised by SRK12 at NT8-4-24 and therefore SCR12 in NT1 is not functional. (C) NT8.4-24 (SRK12) ♀ x NT8.4-24 (SCR12) ♂ = incompatible. Serves as a negative control and confirms that NT8.4-24 accession is an obligate outcrosser and SRK is functional in NT8.4-24. (D) NT8.4-20 (not SRK12) ♀ x NT8.4-24 (SCR12) ♂ = compatible. Serves as a positive control.

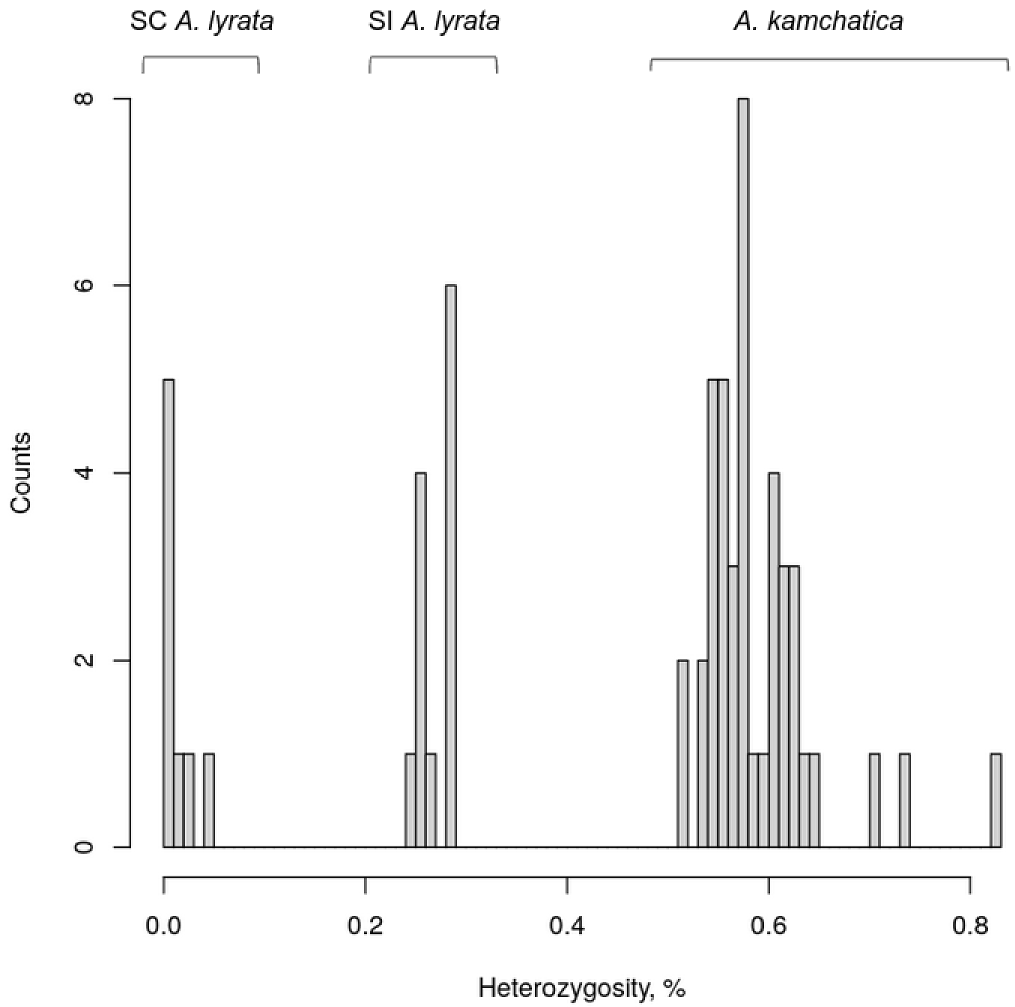

Supplementary figure 12. Three modes of heterozygosity separating self-compatible (SC) A. lyrata, outcrossing or self-incompatible (SI) *A. lyrata* and allotetraploid *A. kamchatica* mapped to NT1 *A. lyrata* reference genome.

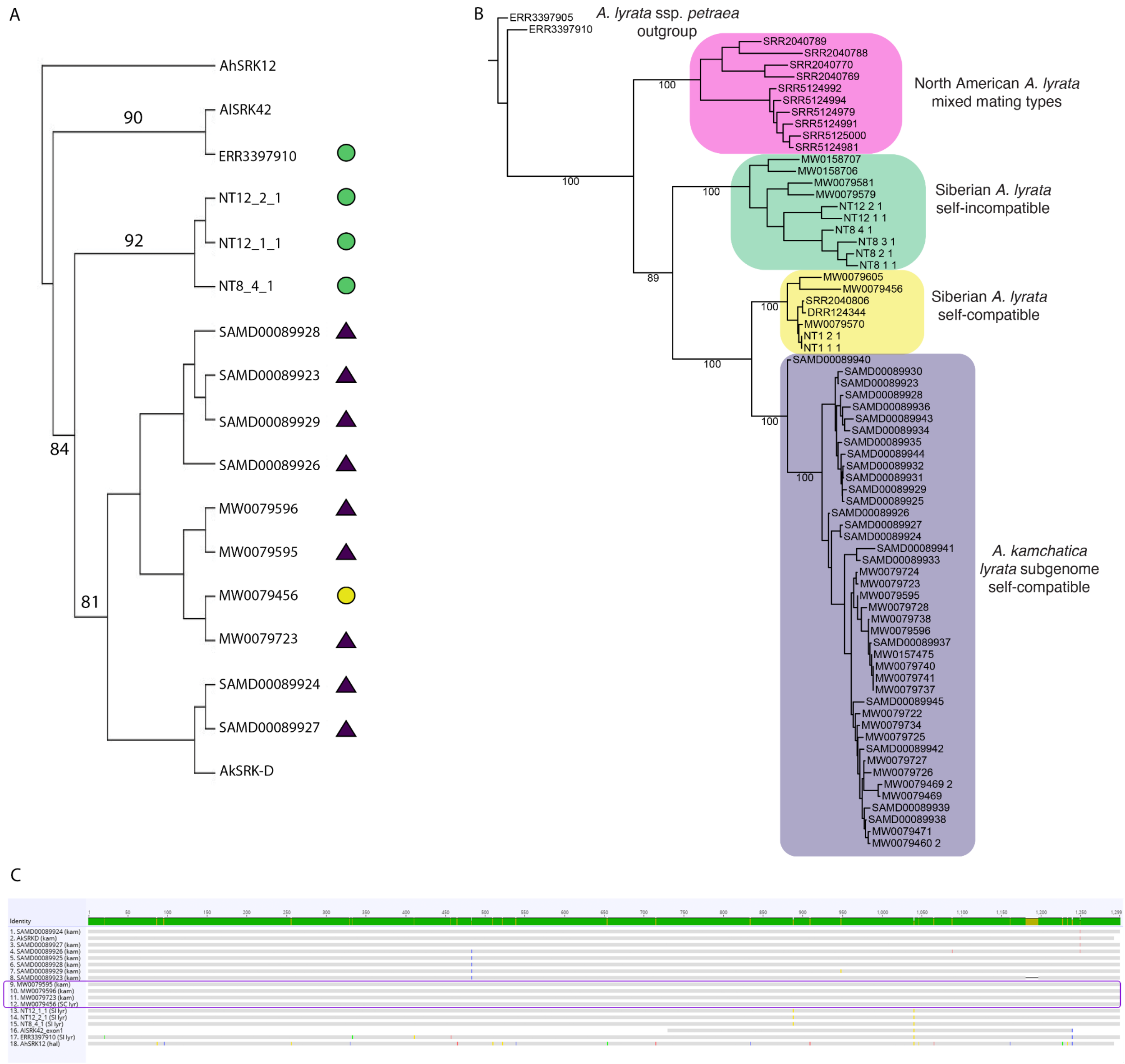

Supplementary Figure 13. (A) Phylogenetic tree with first exons of *SRK* sequences from *A. lyrata* AlSRK42 and *A. kamchatica* (AkSRK-D) rooted on *A. halleri* AhSRK12. Clades with bootstrap support above 80 are labeled. Note that AlSRK42, AkSRK-D and AhSRK12 are members of the same haplogroup, which is older than *Arabidopsis* species. *S*-loci phylogeny between haplogroups does not follow species phylogeny, but it follows the species phylogeny within haplogroups. SRKs of Siberian selfing and outcrossing *A. lyrata* are more closely related to SRK sequences from *A. kamchatica* than to European *A. lyrata* with the same haplogroup. The sequences of AhSRK12 (Goubet et al. 2012), AlSRK42 (Castric et al. 2008) and AkSRK-D (Tsuchimatsu et al. 2012) were published previously. (B) Maximum likelihood tree based on biallelic SNPs shared by the *lyrata* subgenome of *A. kamchatica* and all *A. lyrata* accessions. Bootstrap values only shown for population-level splits, to improve legibility - full Newick tree available as Supplementary Data 4. Clades are colored as in Figure 3, with the addition of North American *A. lyrata* (pink). All *A. kamchatica samples* form a well-supported clade (purple) with self-compatible Siberian *A. lyrata* (yellow). Outside the *A. kamchatica* + selfing Siberian *A. lyrata* clade is self-incompatible Siberian *A. lyrata* (green). The North American clade (pink) includes both self-compatible and self-incompatible accessions, showing that the closest relatives to self-compatible North American *A. lyrata* are outcrossing North American *A. lyrata* and providing further evidence the transition to self-compatibility in Siberia is independent of the transition to self-compatibility in North America. (C) Nucleotide alignment of SRK sequences used for inferring tree in (A). Siberian selfing *A. lyrata* accession MW0079456 has 100% sequence identity with *A. kamchatica* accessions MW0079595, MW0079596, and MW0079723 (purple rectangle).

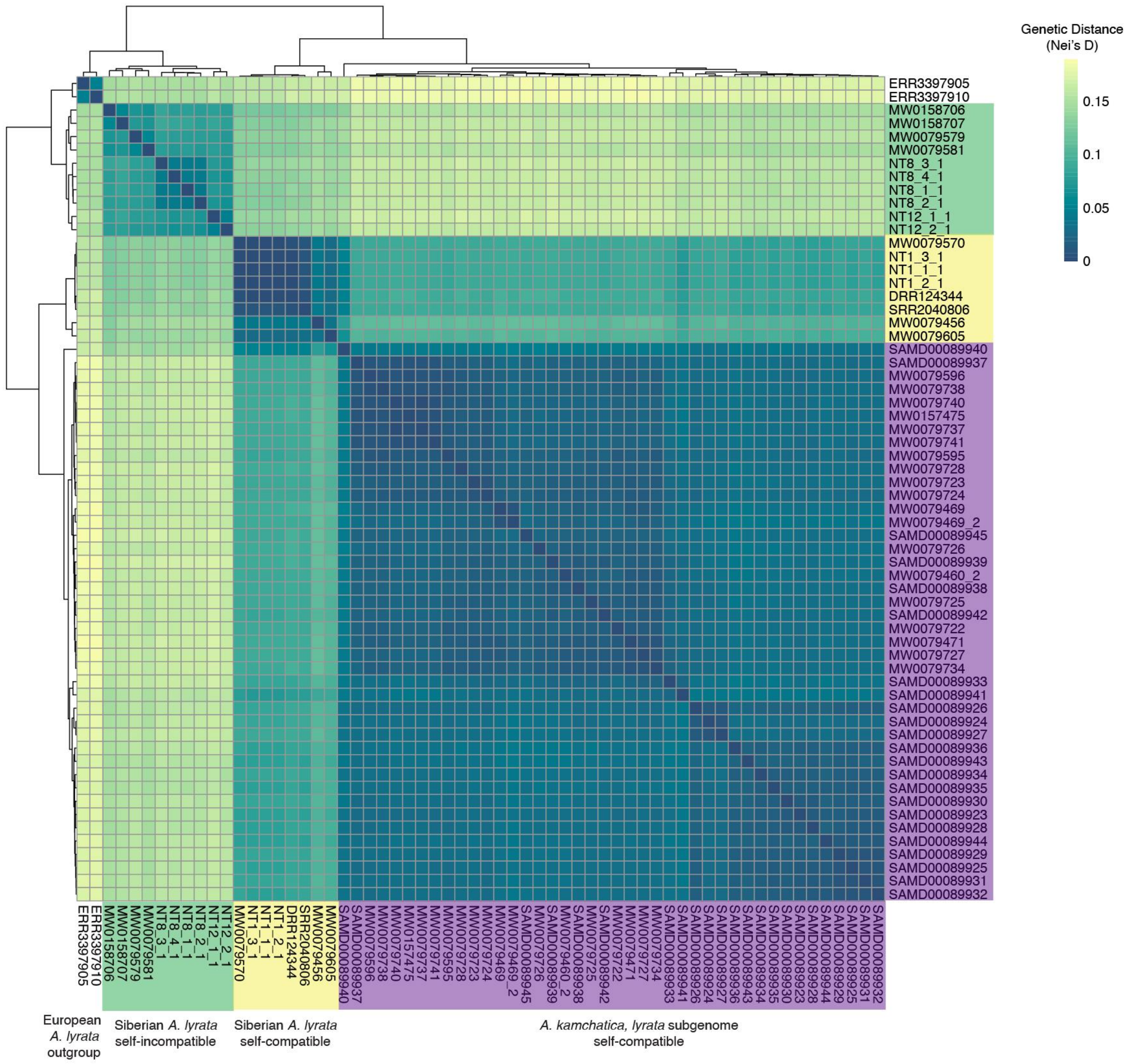

Supplementary Figure 14. Genetic distance (Nei's D) between individual accessions, based on biallelic SNPs and as shown in Figure 3B. Accession names are highlighted to indicate taxon membership - self-incompatible Siberian *A. lyrata* (green), self-compatible Siberian *A. lyrata* (yellow), and *A. kamchatica* (purple). Individual genetic distances among pairs are colored from blue (low distance) to yellow (greatest distance). As a whole, individuals of *A. kamchatica* closer in genetic distance to self-compatible Siberian *A. lyrata* than to outcrossing Siberian *A. lyrata.*

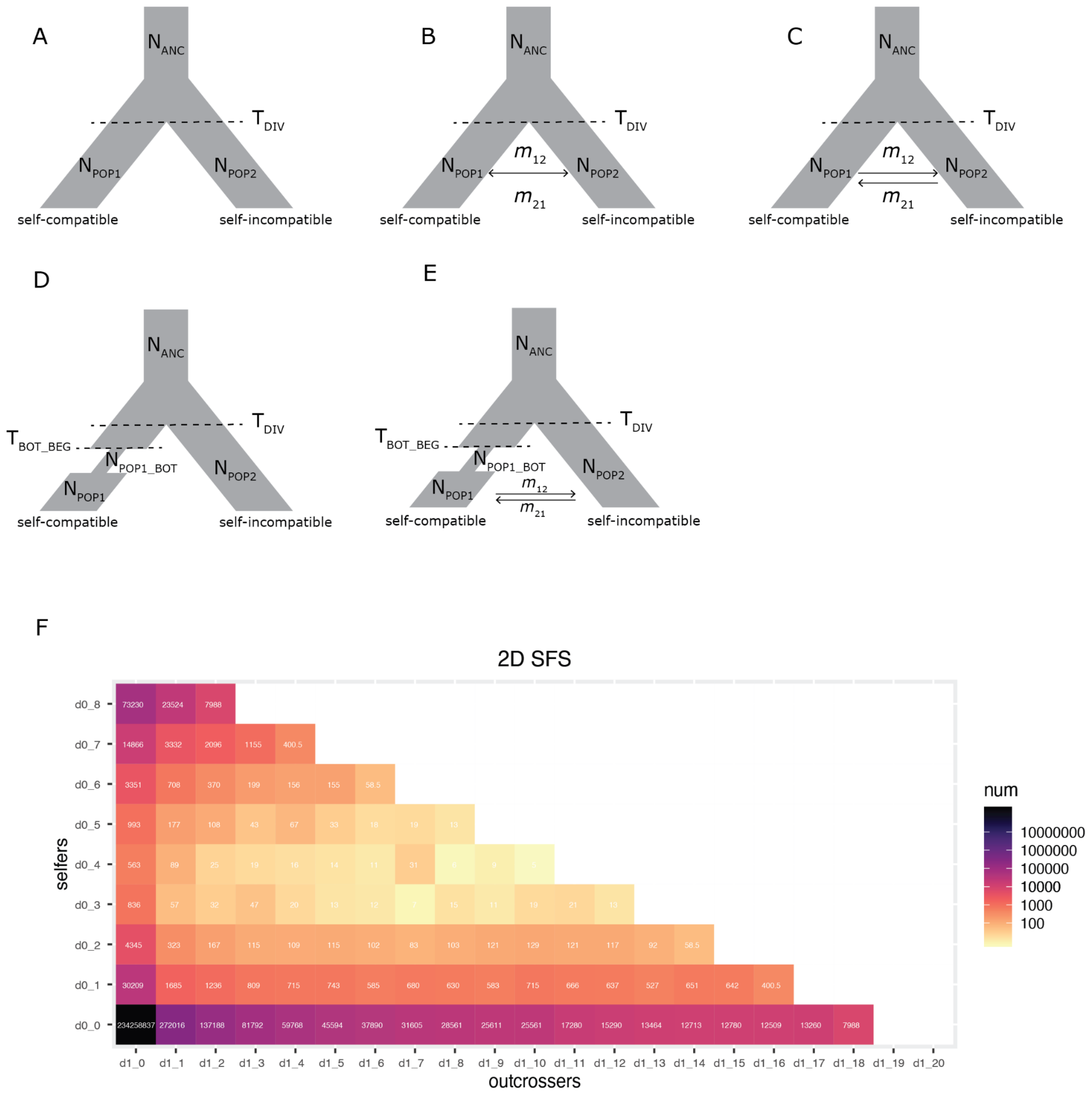

Supplementary Figure 15. Models applied for demographic modeling with fastsimcoal2.
(A) Divergence of self-compatible and self-incompatible populations, without gene flow between them. (B) Model A plus equal bidirectional gene flow (migration) between self-compatible and self-incompatible lineages. (C) Model B but with asymmetric gene flow (migration) between self-compatible and self-incompatible lineages. (D) Simple divergence model as in Model A, plus bottleneck in selfing population. (E) Model C (asymmetric gene flow) plus bottleneck in selfing population. (F) 2D MAF SFS of selfers and outcrossers.

Supplementary Figure 16.

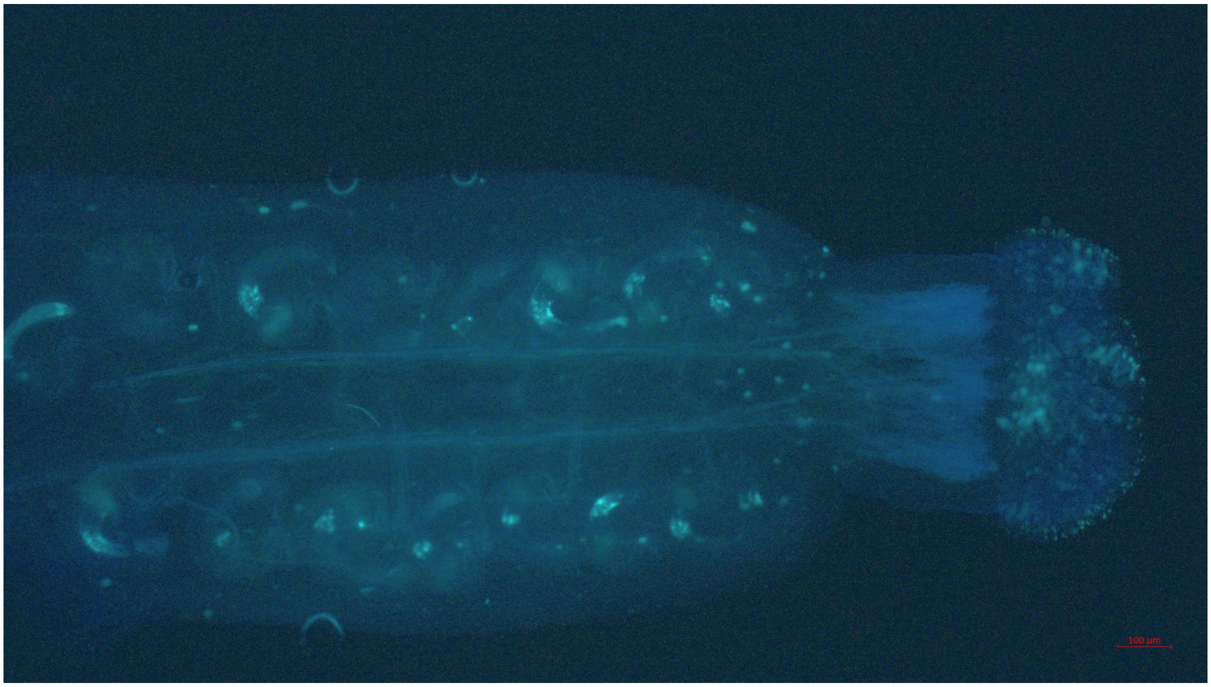

A. TE11.1-2 Selfed. No pollen-tube growth shows self-incompatibility

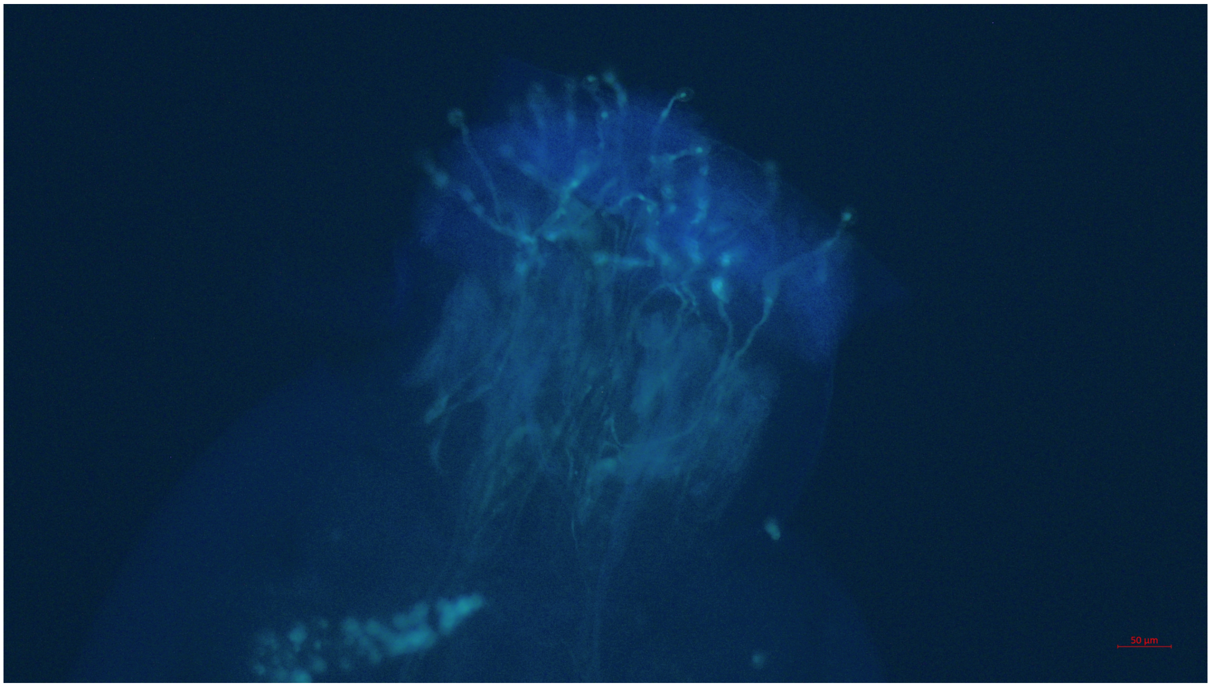

B. TE10.3-2xNT1 F1.1-2 Selfed. Pollen-tube growth shows self-compatibility

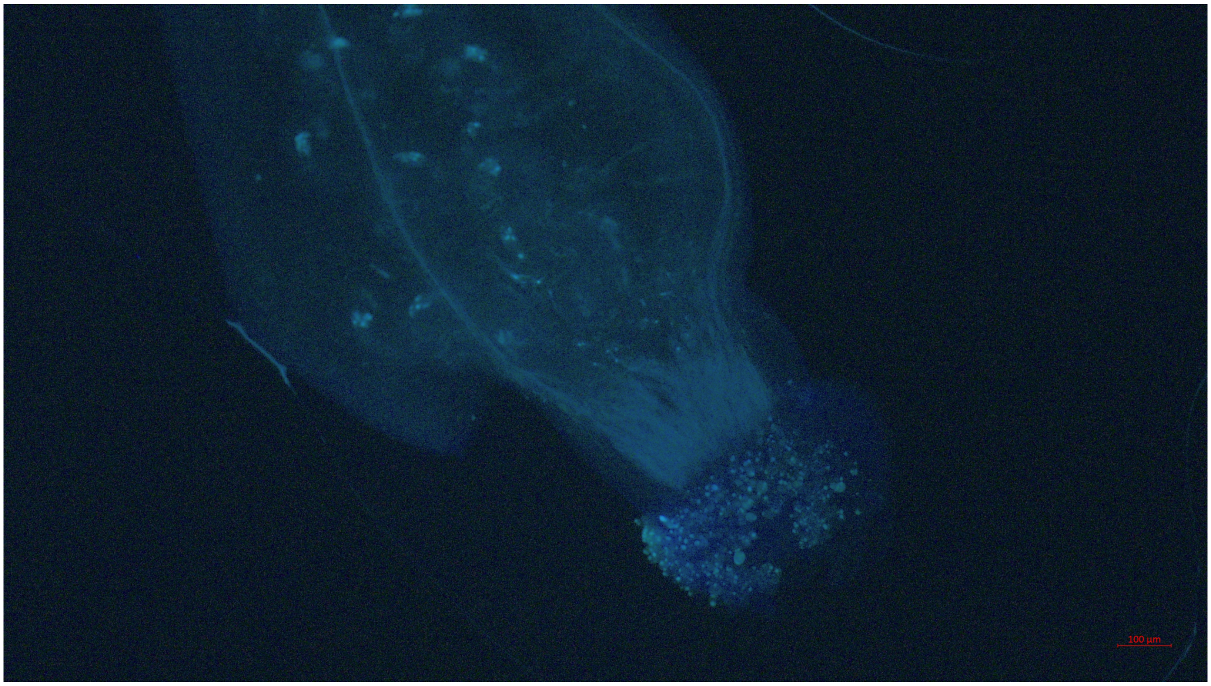

C. TE11.1-2xNT1 F1.2-1 Selfed. No-pollen tube growth

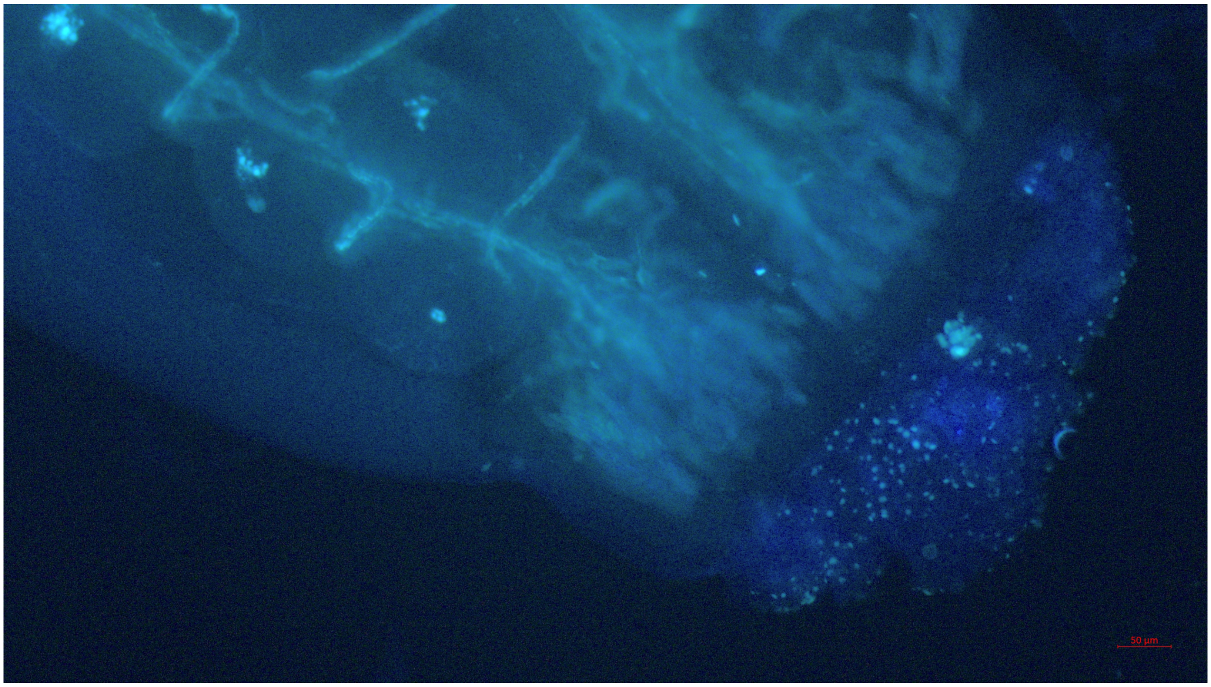

D. TE11-1.2xNT1 F1.2-2 Selfed. No pollen-tube growth shows self-incompatibility
